# Sensing of extracellular L-Proline availability by the integrated stress response determines the outcome of cell competition

**DOI:** 10.1101/2024.11.28.625863

**Authors:** Shruthi Krishnan, Ana Lima, Ying Thong Low, Salvador Perez Montero, Sizhe Tan, Aida Di Gregorio, Adrian Perez Barreto, Sarah Bowling, Karen Vousden, Tristan A. Rodriguez

## Abstract

Cell competition is a quality control acting from development to the adult that eliminates cells that are less-fit than their neighbours. How winner cells induce the elimination of losers during this process is poorly understood. Here, we address this question by studying the onset of differentiation in mouse, where cell competition eliminates 35% of embryonic cells. These loser cells have mitochondrial dysfunction, and we find that this causes amino-acid deprivation and activation of the integrated stress response (ISR), a pathway essential for their survival. We show that L-Proline is a key amino-acid sensed by the ISR and that in a competitive environment, winner cells induce increased L-Proline uptake in loser cells. This causes ISR repression and their elimination. Our results imply that cell competition is acting as a nutrient sensor, eliminating dysfunctional cells when amino acids are plentiful but sparing them in nutrient poor environments.

---

There is an increasing awareness that how cells respond to their environment is shaped not only by their intrinsic genetic composition, but also by the nature of the cells that surround them. One example of this is cell competition, a fitness quality control that results in the elimination of potentially viable cells, but that are less fit than their neighbours^1^. Cell competition has been described from *Drosophila* to humans and been shown to modulate the overall fitness of a tissue in a broad set of contexts^2–6^. These include, for example, the elimination of aberrant cells during embryonic development or ageing, or the expansion of cancerous cells. Interestingly, cell competition not only results in the elimination of defective cells, but in a variation termed super- competition, cells can gain a selective advantage by killing wild-type cells. For example, cells with MYC over-expression^7–9^, loss of function of p53^10–12^ or karyotypic abnormalities^13^ have all been shown to eliminate their wild-type neighbours through competition^14^. A key question is how cells sense their relative fitness levels. However, the mechanisms by which cells with different fitness levels recognise each other is still poorly understood.

To start address this problem, we have studied the competition that occurs at the onset of pluripotent stem cell differentiation. We have shown that during early mouse post-implantation development, about 35% of cells are eliminated through cell competition, and the main features of this elimination can be captured *in vitro* during the differentiation of embryonic stem cells^15–17^. Single cell transcriptomic and functional analysis of the losers in the embryo, identified that these cells showed mitochondrial dysfunction. Furthermore, we also demonstrated that manipulating mitochondrial activity is sufficient to change the competitive ability of pluripotent stem cells^17^, pinpointing mitochondrial function as a key determinant of competitive cell fitness. But how do cells respond to their relative mitochondrial activity?

Mitochondria play a broad range of roles in cellular function, from energy production, the generation of essential metabolites, such as amino acids or nucleotides, and determining the response to nutrient availability in the environment, to the regulation of apoptosis or the REDOX balance of the cell^18, 19^. Consequently, mitochondrial dysfunction can lead to a variety of different stress responses. Interestingly, analysis of the loser epiblast cells in the embryo showed that these presented a signature of activation of the Integrated Stress Response (ISR). The ISR can be activated by diverse stress stimuli, including amino acid deprivation, proteostasis defects, viral infection and Redox imbalance. Once activated, the ISR reduces global translation via phosphorylation of eIF2α, but allows selective translation of selected genes, most notably activated transcription factor 4 (ATF4), that acts to restore cell homeostasis^20, 21^. Here we have investigated the role of the ISR/ATF4 in mouse cell competition and found that it is essential for the survival of cells with mitochondrial dysfunction. We show that in this context, the ISR is activated by amino-acid starvation and promotes amino-acid metabolism in defective cells. Importantly, we find a key role for L-Proline and identify that when cells with mitochondrial dysfunction are surrounded by wild-type cells, they are unable to sustain ISR activation due to the uptake of extracellular L-Proline, that induces their elimination. These results indicate that in a heterogeneous cell population, wild-type and stressed cells respond to the microenvironment differently, and this difference affects tissue composition by deterring if stressed cells will survive or die.

## Results

### Loser cells show activation of the ISR

Here we set out to determine how winner and loser cells communicate their fitness levels during cell competition occurring in the early mouse embryo, where we have previously shown it to be responsible for the elimination of about 35% of epiblast cells^16, 17^. Three possible modes of competition have been proposed, competition for nutrients, where loser cells are less able to uptake essential nutrients, mechanical competition, where loser cells are eliminated at high densities due to their increased sensitivity to compaction, and fitness sensing, where winners and losers sense their relative fitness levels^6^. To distinguish between these possibilities, we first studied the mode of elimination of mouse embryonic stem cells (ESCs) with defective BMP signalling (*Bmpr1a^-/^*^-^) co-cultured with wild-type cells, as it recapitulates the most important features of the competition occurring in the embryo^15–17^. Wild-type and mutant cells were mixed in a 1:1 ratio and plated at increasing densities. Cells were then induced to differentiate for 4 days by culture in N2B27^15^. Interestingly, we found that although BMP signalling defective ESCs were effectively eliminated at low densities, this elimination was inhibited as cell density increased (Supp. Fig.1). The decreased competitive elimination of defective cells at high densities suggests that neither mechanical force nor competition for nutrients are likely to be determining the outcome of cell competition.

Given that plausibility of the fitness sensing model, we hypothesized that one mechanism by which winner and loser cells could sense each other is through the differential activation of stress pathways. We have previously shown that a key feature of loser cells is their mitochondrial dysfunction^17^. In the embryo, the cells eliminated by competition additionally display a transcriptional signature of ISR activation^17^, a pathway downstream of several stresses associated with mitochondrial dysfunction, including amino acid deprivation, ER stress and Redox imbalance. To validate these results, we cultured E5.5 embryos in the presence a pan-caspase inhibitor (CI) or DMSO-vehicle control for 18h (Fig.1A). After treatment, the embryos were stained for the ISR markers, ATF4 and CHOP. We found both to be increased in the epiblast cells of CI-treated as compared to DMSO-treated control embryos (Fig.1B-C). This supports that loser cells in the embryo exhibit ISR activation.

**Figure 1.**
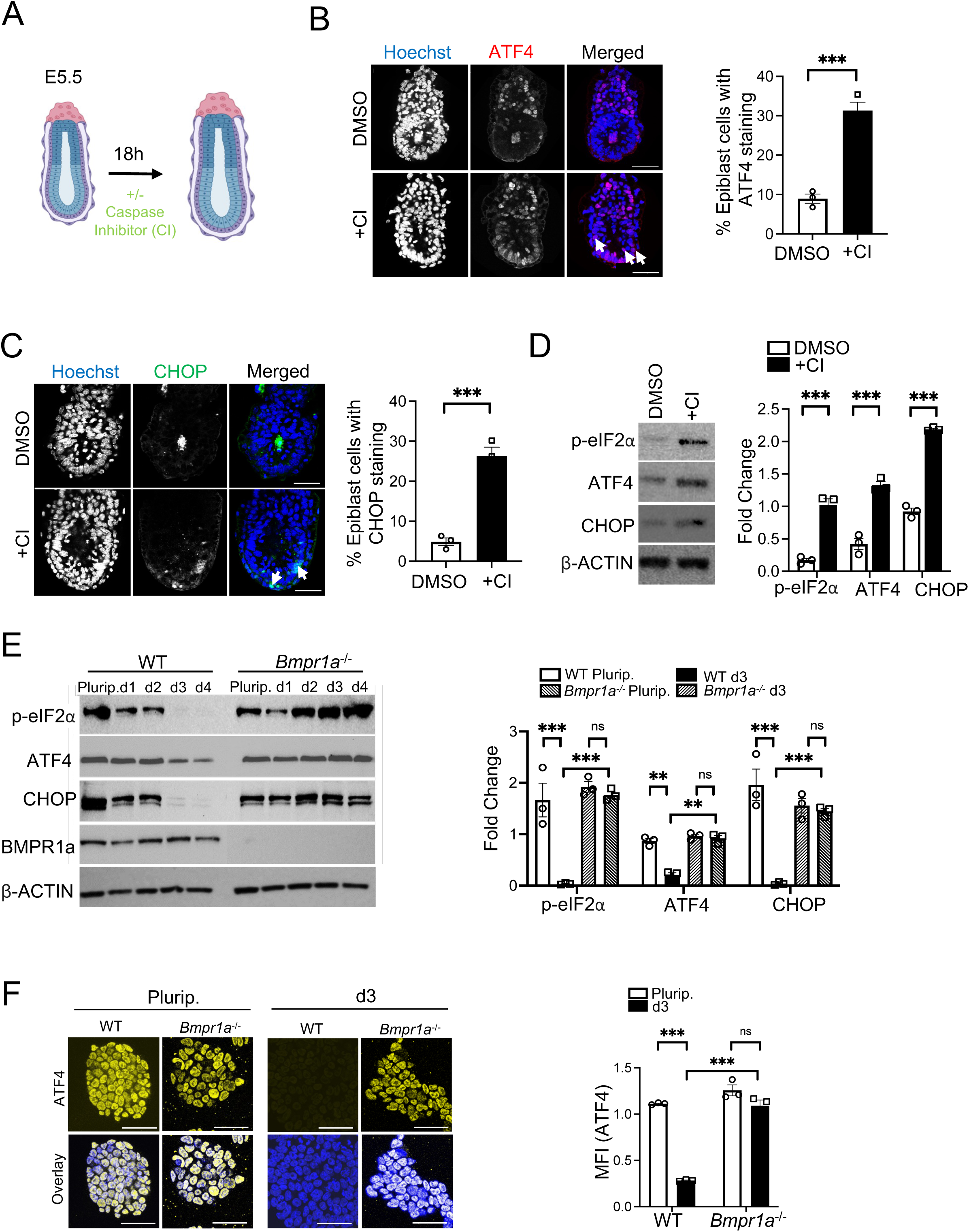
Defective cells eliminated by cell competition have a signature of ISR activation. A, Experimental design. E5.5 embryos were treated with or without the pan caspase inhibitor, Z-VAD-FMK (CI; 100uM) for 18h. B-C, Immunofluorescence analysis of ATF4 (red; B) and CHOP (green; C) in E6.5 embryos treated with CI or DMSO vehicle control for 18h. Embryos were counterstained with Hoechst (blue). Scale bars= 50uM. Bar graphs depict quantification of epiblast cells stained positive for ATF4 (B) and CHOP (C). D, Wildtype ESCs were treated at day 2 of differentiation with CI (100uM) or DMSO for 24h and immunoblots were performed to analyse the levels of ISR proteins; phosphorylated eIF2α (p-eIF2α), ATF4 and CHOP. β-ACTIN was used as a loading control. Bar graph represents densitometry-based quantification of fold change in expression of each the proteins as a value normalised to the respective loading control. E, Immunoblot analysis of wildtype (WT) and *Bmpr1a^-/-^* cells after culture in pluripotency conditions (Plurip.) or at different time-points of differentiation (d1-d4). Expression of p-eIF2α, ATF4, CHOP BMPR1a and β-ACTIN (loading control) is analysed. Bar graphs represent fold change in expression levels of p-eIF2α, ATF4 and CHOP in ESC and d3. F, Immunofluorescence analysis of ATF4 (yellow) in WT and *Bmpr1a^-/-^* ESCs cultured in pluripotency conditions or at day 3 of differentiation. Nuclei are counterstained with Hoechst (blue). Scale bars= 100uM. Bar graph represents quantification of staining intensity of ATF4 (MFI in arbitrary units) in the cells. n=3 for all studies. Error bars denote SEM. *** p < 0.005; unpaired, t-test (B-D) or two-way ANOVA and Tukeys post-hoc test (E, F).

We next tested if the ISR is activated in the cells dying during the first steps of ESC differentiation, as this models early post-implantation epiblast development^15^. We differentiated wild-type ESCs for 3 days in N2B27, treated them with CI or DMSO for 24h and assessed p-eIF2α, ATF4 and CHOP expression. We found increased levels of expression for all three ISR markers in CI-treated cells when compared to DMSO-treated controls (Fig.1D), suggesting that the ISR is also marking the cells eliminated during differentiation. This led us to ask if there also is a differential ISR response in the defective ESCs eliminated by cell competition. For this we studied two loser cell models with mitochondrial dysfunction^17^, the above mentioned *Bmpr1a^-/-^* ESCs, and *Drp1^-/-^*cells, that carry a null mutation in the regulator of mitochondrial fission, *Drp1*. In agreement with the high expression of p-eIF2α and ATF4 in the pre-implantation embryo and ESCs^22^, we found that wild- type and mutant cells display activation of ISR markers when cultured as homogeneous populations in pluripotency conditions (Fig.1E-F and Supp. Fig.2A-B). In contrast to this, from day 3 of differentiation, we found that whilst wild-type cells strongly downregulate ISR protein expression, both defective cell types failed to do so and sustained robust activation of this pathway (Fig.1E-F and Supp. Fig.2A-B). The fact that we only see competition from day 3 of differentiation^15–17^, suggests that this differential ISR activation between winner and loser cells may regulate their competitive behaviour.

### Activation of the ISR is essential for loser cell survival during cell competition

To establish the role of the ISR pathway during cell competition, we first analysed the expression of ATF4 and CHOP in co-cultured wild-type and *Bmpr1a^-/-^*ESCs. Immunofluorescence analysis revealed that whilst in separate culture the expression of ATF4 and CHOP was high in mutant cells and low in wild-type cells at day 3 of differentiation, this situation was reversed in co-culture. In this condition, the *Bmpr1a^-/-^* cells showed a downregulation of ATF4 and CHOP expression, and the wild-type cells upregulated the expression of these factors (Fig.2A-B). The decreased ISR expression found in *Bmpr1a^-/-^*cells in co-culture suggests that this pathway may be regulating loser cell survival.

**Figure 2.**
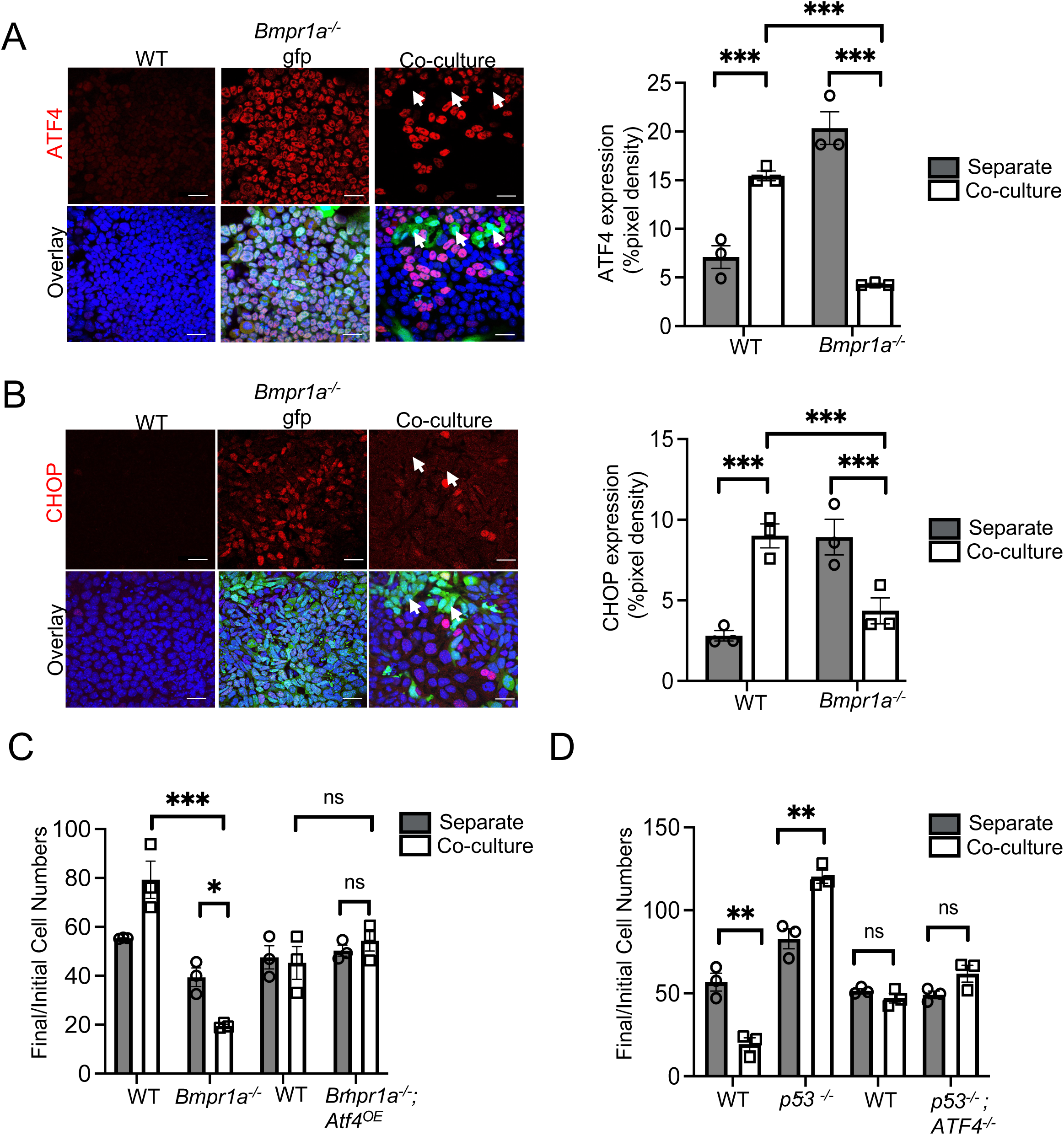
Activation of ISR is essential for loser cell elimination. A, B, Immunofluorescence analysis of ATF4 and CHOP (red) in wild-type (WT) and *Bmpr1a^-/-^*-Gfp cells cultured in separate and co-culture conditions and with the nuclei counterstained with Hoechst (blue); white arrows depict lack of ATF4 and CHOP staining in Gfp-tagged *Bmpr1a^-/-^* cells and white arrow heads depict colocalization of ATF4 and CHOP staining in WT cells. Scale bars= 50uM for staining in separate cultures of WT and *Bmpr1a^-/-^*-Gfp cells and scale bars=100uM for staining in co-cultures. Bar graphs depict ATF4 (A) and CHOP (B) expression as % of pixel density in the images. C, Cell competition assays between WT and *Bmpr1a^-/-^* cells as well as between WT and *Bmpr1a^-/-^*; *Atf4* OE (OE; overexpressing) cells. Bar graph represents ratio between final (day 4) cell numbers to initial cell numbers in separate and co-cultures for each of these cell types. D, Bar graph depicts ratio between final (day 4) cell numbers to initial cell numbers of WT, *p53^-/-^* and WT, *p53^-/-^*; *Atf4^-/-^* cells in separate and co-cultures. n=3 for all studies. Error bars denote SEM. *** p < 0.005, ns-non-significant; two-way ANOVA and Tukeys post-hoc test (A-D).

To test the above possibility, we did two things. First, we analysed the impact of *Atf4* loss of function in ESCs and during differentiation to understand the effects of downregulating this factor. We found that although *Atf4^-/-^* cells displayed normal proliferation when cultured in pluripotency conditions, their growth was severely compromised when induced to differentiate (Supp. Fig.2C- D). This diminished proliferation was primarily due to apoptosis, as mutant cells cultured in N2B27 showed a marked increase in cleaved caspase 3 expression (Supp. Fig.2E), but no change in their cell cycle length, as inferred by the rate of EdU incorporation (Supp. Fig.2F). We next over- expressed *Atf4* in *Bmpr1a^-/-^* cells under the control of a ubiquitous promoter (*Bmpr1a^-/-^; Atf4^OE^*). Notably, we found that this was sufficient to completely rescue the elimination of mutant cells in co-culture (Fig.2C). This indicates that the ATF4 downregulation observed in *Bmpr1a^-/-^* cells in co-culture is likely causing their elimination.

Given that we had also found an upregulation of ATF4 in co-cultured wild-type cells (Fig2.A-B), we tested the importance of ATF4 for winner cell behaviour. For this, we resorted to a super-competition model, where *p53^-/-^* eliminate wild-type cells^10^ and mutated *Atf4* in the *p53^-/-^* cells. We found that *p53^-/-^; Atf4^-/-^* cells no longer have a competitive advantage over wild-type cells, as they are no longer able induce their elimination (Fig.2D and Supp. Fig.2G). These results highlight the importance of ATF4 for determining how stressed and unstressed cells adapt to a heterogeneous environment. In the stressed cells ATF4 downregulation is driving their elimination and in the unstressed cells ATF4 upregulation is supporting their competitive advantage. But the question that then arises is how does ATF4 play these roles?

### Amino acid starvation activates the ISR to promote amino acid biosynthesis

To address the mechanism by which ATF4 determines the outcome of cell competition, we performed transcriptomic comparisons of separate cultures of *Atf4^-/-^* versus wild-type cells and of *p53^-/-^; Atf4^-/-^* versus *p53^-/-^* cells at day 3 of differentiation. Differential cellular pathway analysis by KEGG enrichment revealed that the topmost downregulated pathways in both sets of cells with *Atf4* mutations were those associated with non-essential amino acid biosynthesis, transport and metabolism (Fig.3A-B). This agrees with the finding that the ISR reroutes carbon utilization away from the tricarboxylic acid cycle and towards amino acid production^23^. To further examine this possibility, we differentiated *Atf4^-/-^* ESCs in the presence of excess non-essential amino acids and found that this partially rescued the growth defect of mutant cells (Supp. Fig.2H). Testing individual or combinations of amino acids revealed that Proline and Asparagine could recapitulate the rescue seen with the pool of non-essential amino acids (Supp. Fig. 2H). These results support that the ISR is determining the outcome of cell competition by regulating amino acid levels.

**Figure 3.**
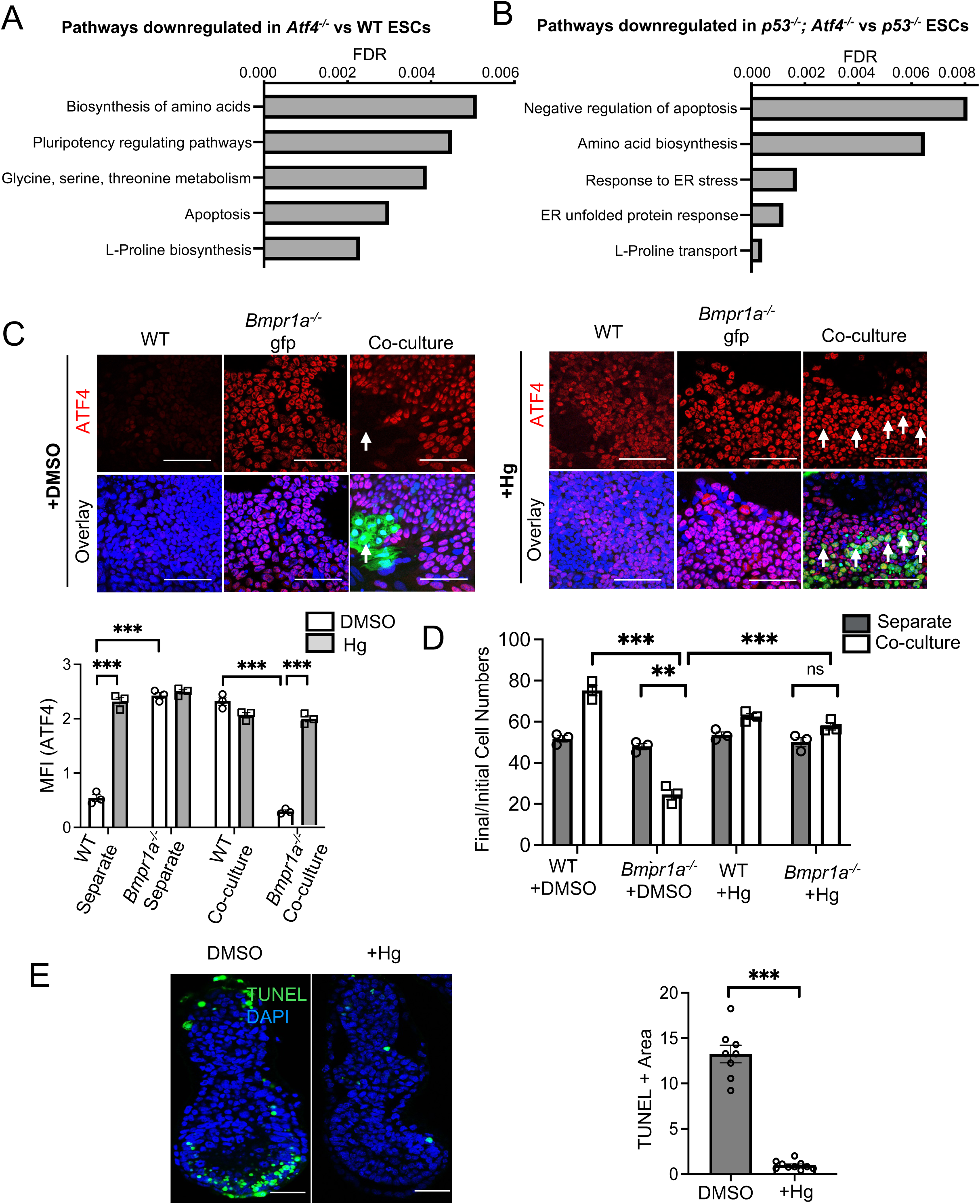
Activation of the amino-acid depravation branch of the ISR prevents cell competition. A, B, Top downregulated pathways identified by KEGG enrichment analysis of the transcriptomic data sets obtained from *Atf4^-/-^* vs wild-type (WT) cells (A) and p53^-/-^; *Atf4^-/-^* vs p53^-/-^ cells (B). C, Immunostaining analysis for ATF4 (red) in separate and co-cultures of WT and *Bmpr1a^-/-^-*Gfp cells cultured for 48h in DMSO or Halofuginone (Hg; 20nM) containing media. Nuclei are counterstained with Hoechst (blue). Scale bars= 100uM. Bar graph represents quantification of staining intensity of ATF4 (MFI in arbitrary units) in these cells. D, Cell competition assays between WT and *Bmpr1a^-/-^* cells cultured for 48h in DMSO or Hg treated media depicted as the ratio of final (day 3) cell numbers to the initial cell numbers of the cells in separate and co-cultures. E, Embryos (E5.5) were cultured for 18h in medium supplemented with Hg (20nM) or DMSO. TUNEL assay (green) was performed to assess cell death in the epiblast cells of these embryos. Nuclei were counterstained with DAPI (blue). Scale bars=50uM. Bar graph represents epiblast area stained positive for TUNEL as an indicator of cell death in epiblasts of the embryos. n=3 for A-D, n=8 for B. Error bars denote SEM. *** p < 0.005, **p < 0.01, ns-non-significant; two-way ANOVA and Tukeys post-hoc test (C, D). *** p < 0.005, unpaired t-test (E).

One of the key drivers of ISR activation is amino acid starvation, as amino acid depletion triggers the sensor kinase GCN2 to phosphorylate eIF2α and activate the ATF4-CHOP axis^20, 21^. Given our finding that ATF4 primarily regulates amino acid biosynthesis in differentiating ESCs, one possibility is that defective cells activate the ISR in response to an amino acid deficiency caused by mitochondrial dysfunction. In this scenario, the downregulation of the ISR in co-culture could be due to these cells no longer sensing a lack of amino acids. Halofuginone is a glutamyl-prolyl tRNA synthetase inhibitor that induces accumulation of uncharged prolyl tRNAs and activation of the amino acid starvation response^24^. To test the above hypothesis, we differentiated wild-type, *Bmpr1a^-/-^* and *Drp1^-/-^* cells separately or as wild-type/defective cocultures and treated them for 48 hours with Halofuginone. We found that this increased ATF4 expression in all cell types and restored expression in *Bmpr1a^-/-^* and *Drp1^-/-^* cells to wild-type levels, both in separate and co-culture conditions (Fig.3C and Supp. Fig.3A-B). Importantly, this increase in ATF4 was sufficient to rescue defective cell elimination in co-culture (Fig.3D and Supp. Fig.3C). In the embryo, cell competition leads to a wave of cell death at E6.0^7, 15, 16^. We therefore examined the importance of activation of the ISR for this cell death, by culturing E5.5 embryos for 18 hours in the presence of Halofuginone. We found that this abolished the cell death occurring at this stage in the embryo (Fig.3E). It is also noteworthy that a proportion of Halofuginone treated embryos presented epiblast malformations, suggesting that the amino acid starvation response may also be regulating post-implantation embryo morphogenesis. Together, these results indicate that activation of the amino acid starvation branch of the ISR with Halofuginone can rescue loser cell elimination during cell competition. This points to repression of the amino acid starvation branch of the ISR as a key step driving the elimination of defective cells in a competitive environment.

### Extracellular L-Proline determine the outcome of cell competition

The finding that defective cells no longer sense amino acid deprivation in co-culture, suggests that in this condition they are exposed to increased levels of one or several amino acids. To test this possibility, we first set out to establish which amino acids repressed the ISR during cell competition. Our transcriptional profiling of Atf4 mutant cells suggested that this factor regulates L-Proline biosynthesis and transport (Fig.3 and Supp. Fig.2H), an amino acid previously shown to repress ATF4 in ESCs^25^. We therefore first analysed the ability of L-Proline to repress the ISR by treating wild-type, *Bmpr1a^-/-^* and *Drp1^-/-^* ESCs cultured separately in pluripotency and differentiation conditions for 48 hours with excess L-Proline. We found that L-Proline induced a downregulation of p-eIF2α, ATF4 and CHOP in in all cell types and in all conditions analysed (Fig.4A-B and Supp. Fig.4A-B). Furthermore, we also observed that L-Proline treatment induced cell death, as measured by cleaved CASPASE 8 levels, in wild-type, *Bmpr1a^-/-^* and *Drp1^-/-^* cells when induced to differentiate (Supp. Fig.4C). These results suggest that excess L-Proline may be inducing the ISR downregulation that causes defective cell elimination in co-culture.

**Figure 4.**
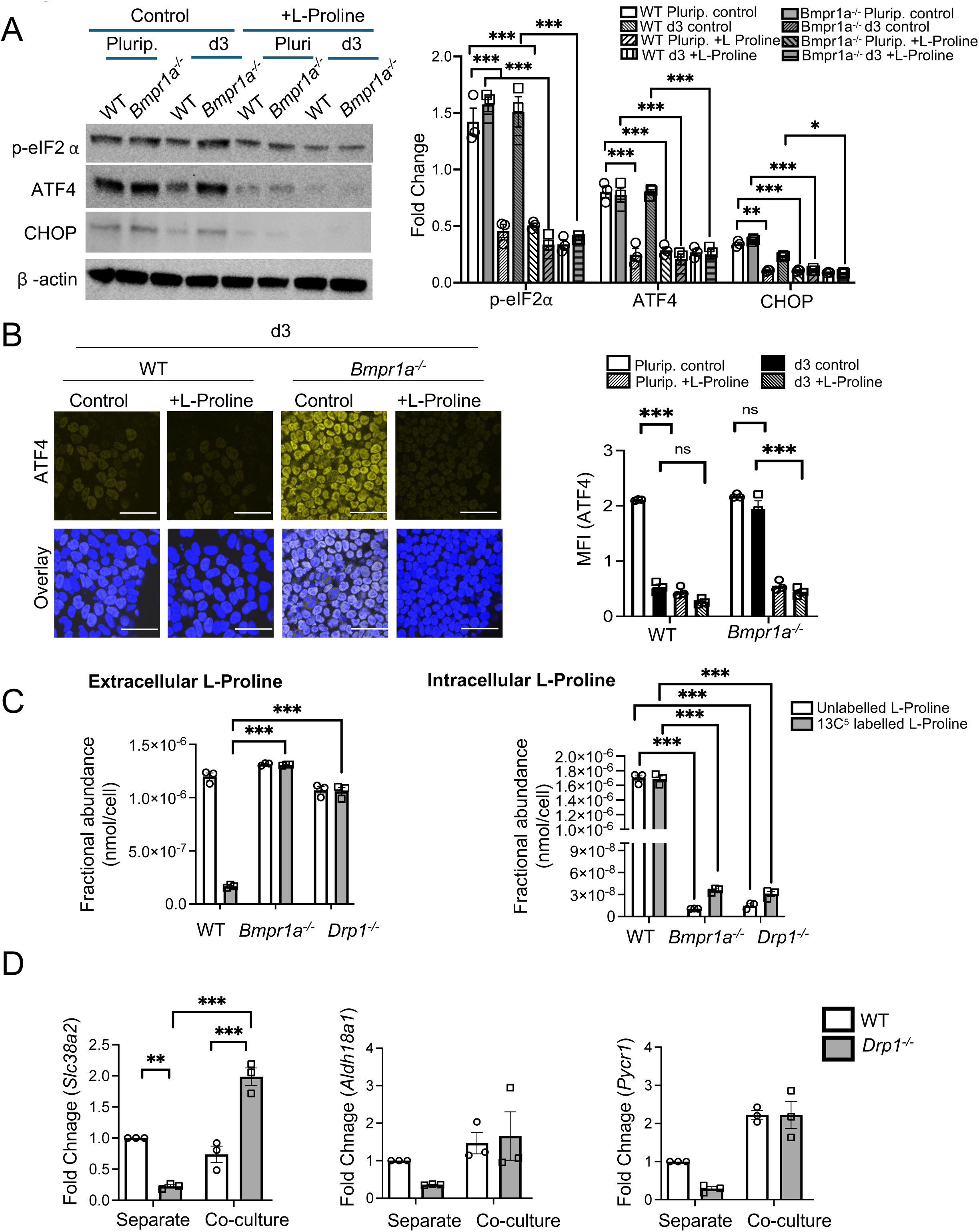
L-Proline represses ATF4-mediated ISR signaling. A, Wild-type (WT) and *Bmpr1a^-/-^*ESCs were cultured in pluripotency conditions (Plurip.) or induced to differentiate for 3 days (d3) cultured in control medium or medium supplemented with L-Proline for 48h (0.5mM; +L-Proline) and the expression of ISR proteins, p-eIF2α, ATF4, CHOP was analysed by immunoblots. β-Actin; loading control. Bar graph represents fold change in expression of the proteins in the various conditions. B, Immunofluorescence analysis of ATF4 in WT and *Bmpr1a^-/-^* ESCs at day 3 of differentiation cultured in control medium or medium supplemented with L-Proline for 48h. Scale bar=100uM. Bar graph represents quantification of staining intensity of ATF4. C, Bar graphs represent fractional abundance of extracellular and intracellular L-Proline obtained by GC-MS of the media samples and cell extracts from WT, *Bmpr1a^-/-^* and *Drp1^-/-^* cells grown as homogenous population at day 3 of differentiation. D, Quantitative RT-PCR showing expression levels of the gene encoding neutral amino acid transporter responsible for L-Proline uptake (*Slc38a2*) and of the genes encoding the enzymes that regulate L-Proline synthesis (*Aldh18a1 and Pycr1*) in separately and co-cultured WT and *Drp1^-/-^* ESCs at day 3 of differentiation. Gene expression is normalized against *Gapdh*. n=3 for all studies. Error bars denote SEM. *** p < 0.005, **p < 0.01, *p < 0.05, ns-non-significant; two-way ANOVA and Tukeys post-hoc test.

Next, we analysed L-Proline metabolism in wild-type, *Bmpr1a^-/-^* and *Drp1^-/-^* cells. For this we first separately cultured these cells in L-Proline deficient medium supplemented with labelled L-Proline (^13^C_5_-L-Proline). We replaced the cells with fresh labelled L-Proline-containing medium every 24 hours from day 0 of differentiation and collected both the medium and cells for metabolite extraction at day 3 of differentiation. This allows to analyse L-Proline synthesis, by measuring the presence of un-labelled L-Proline, as well as its uptake, by studying the extra-cellular versus intracellular localization of ^13^C_5_labelled L-Proline. GC-MS analysis of supernatants indicated that at day 3 of differentiation both wild-type and defective cells had similar levels of extra-cellular un- labelled L-Proline, indicating that the three cell types had a similar capacity to secrete this amino-acid. In contrast to this, when we analysed labelled L-Proline levels, we found that whilst *Bmpr1a^- /-^* and *Drp1^-/-^*supernatants showed higher L-Proline levels than wild-type cells, their cell pellets displayed significantly lower levels (Fig.4C). The high extra-cellular and low intracellular levels of L-Proline in *Bmpr1a^-/-^* and *Drp1^-/-^* cells indicates that these cells have a deficiency in L-Proline uptake. When we analysed the levels of un-labelled L-Proline in cell pellets, we found that mutant pellets also had lower levels than those from wild-type cells (Figure4C). The lower levels of intracellular, newly synthesized/un-labelled L-Proline, signifies that *Bmpr1a^-/-^* and *Drp1^-/-^* cells also likely have defective L-Proline synthesis. Given that L-Proline represses ATF4 (Fig.4A-B and Supp. Fig.4A-B), the lower L-Proline uptake and synthesis observed in mutant cells explains the elevated ATF4 expression seen in these cells in separate culture at day 3 of differentiation (Fig.1).

To understand the causes for the differing ability of wild-type and mutant cells to uptake and synthesize L-Proline, we analysed in wild-type and *Drp1^-/-^* cells the expression levels of *Sclc38a2*, the gene that encodes the neutral amino acid transporter responsible for L-Proline uptake and of *Aldh18a1* (encoding P5CS) and *Pycr1*, that regulate de novo L-Proline synthesis. In agreement with the lower intra-cellular levels of L-Proline observed in defective cells, we found that in separate culture *Drp1^-/-^*cells show a downregulation in the expression of both the genes responsible for L-Proline uptake as well as for its synthesis (Fig.4D). In contrast to this, when we analysed the expression of these factors in co-culture, we found that *Aldh18a1* and *Pycr1* were up-regulated in mutant cells to the levels found in wild-type cells, and *Sclc38a2* was increased in *Drp1^-/-^* cells to 3-fold higher levels than those of wild-types (Fig.4D). This suggest that the presence of wild-type cells have induced mutant cells to increase L-Proline uptake and synthesis. This finding correlates with the repression of ATF4 expression that we see in mutant cells in co-culture (Fig.2) and provides a likely explanation for their elimination.

The above results suggest that the uptake of extracellular L-Proline may determine the outcome of cell competition, with the increased ability to do so of defective cells in co-culture compared to separate culture driving their elimination. To test this possibility, we performed our cell competition assays in media without L-Proline (-L-Proline). When either *Bmpr1a^-/-^* or *Drp1^-/-^* cells were co-cultured with wild-type cells in L-Proline depleted media, we found that the lack of exogenous L-Proline led to a marked increase in ATF4 expression in both defective cell types (Fig.5A and Supp. Fig.5A-B). Notably, the levels of ATF4 expression in co-cultured defective cells were now equivalent to those found in these cells in separate cultures. L-Proline deficient media also increased ATF4 expression in wild-type cells in separate culture but not in co-culture, meaning that in this condition, wild-type and defective cells now have the same levels of ATF4 expression. The increase in ATF4 expression in defective cells was accompanied by a decrease in apoptosis, as measured by the levels of cleaved-caspase 8 (Fig.5B and Supp. Fig.6A-B). Importantly, the co-culture of defective cells with wild-type cells in media lacking L-Proline completely rescued their elimination (Fig.5C-D). Notably, this rescue was not due to a block in differentiation, as L-Proline deficient media did not impair the ability of ESCs to differentiate (Supp. Fig.7A) and adding back half the normal amount of L-Proline from day 1 of differentiation restored the competitive ability of the cells (Supp. Fig.7B-C).

**Figure 5.**
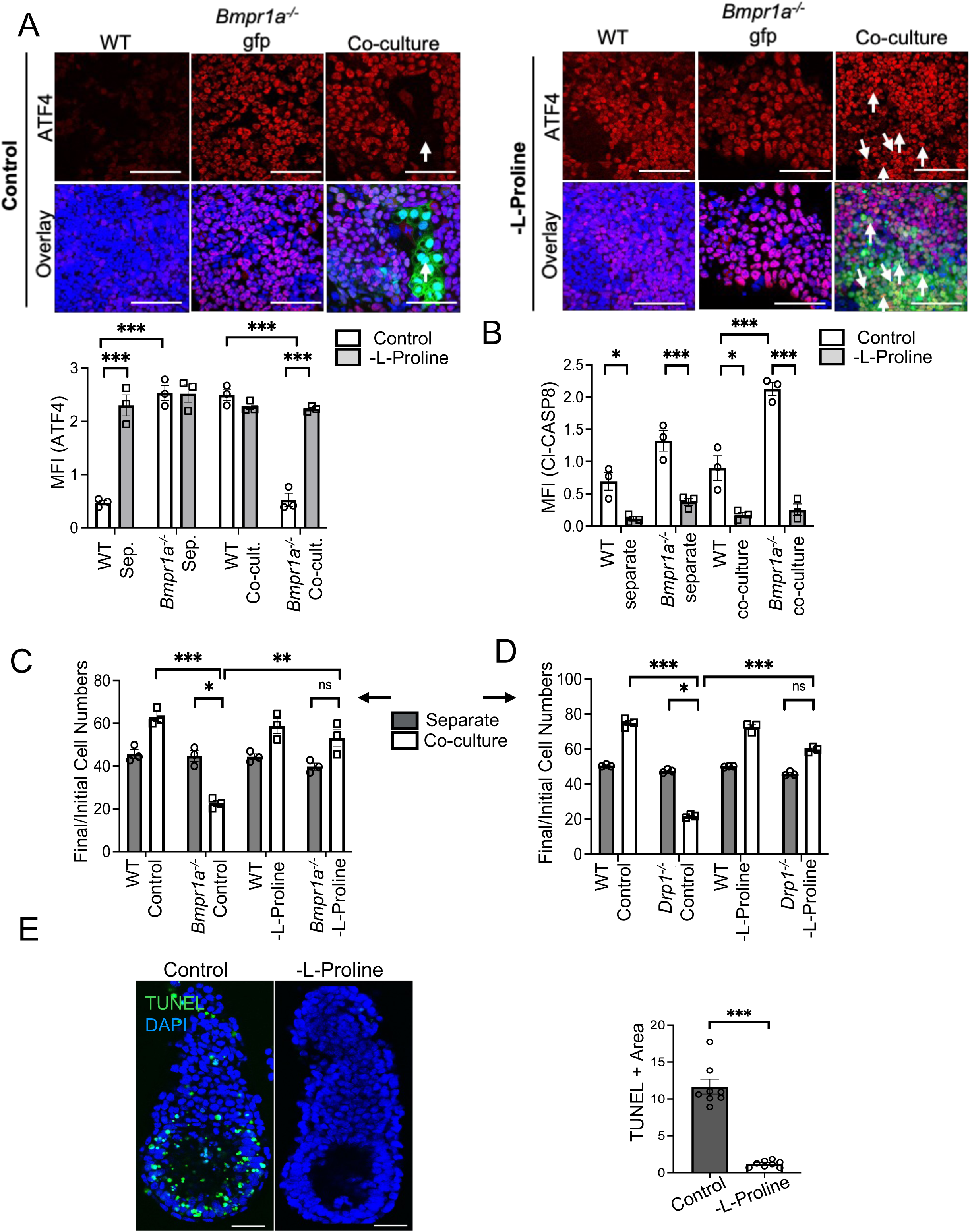
L-Proline mediated regulation of ATF4 levels leads to competitive loser cell elimination. A, Immunofluorescence analysis for ATF4 (red; C) in separate and co-cultures of wild-type (WT), and *Bmpr1a^-/-^* cells, cultured in control and L-Proline deficient medium (-L-Proline media). Nuclei is counterstained with Hoechst (blue). Scale bars=100uM. Bar graphs represent quantification of staining intensity of ATF4. B, Bar graph represents quantification of staining intensity of cleaved caspase 8 in separate and co-cultures of WT, *Bmpr1a^-/-^* cells, cultured in control and -L-Proline media. C, D, Bar graphs representing ratio of final (day 3) to initial cell numbers of WT, *Bmpr1a^-/-^* (C) and WT, *Drp1^-/-^* cells (D) in control and -L-Proline medium for separate and co-cultures. E, Embryos (E5.5) were cultured for 18h in control (N2B27) and -L-Proline medium. TUNEL assay (green) was performed to assess cell death in epiblast cells of the embryos, nuclei counterstained with DAPI (blue). Scale bars=50uM. Bar graph represents epiblast area stained positive for TUNEL as an indicator of cell death in epiblasts of the embryos. n=3 for A-D, n=8 for E. Error bars denote SEM. *** p < 0.005, **p < 0.01, *p < 0.05, ns-non-significant; two-way ANOVA and Tukeys post-hoc test.

We next tested the in vivo relevance of this observation. When E5.5 embryos were cultured in media lacking L-Proline for 18 hours, we found that this was sufficient to abolish the cell death occurring in the epiblast (Fig.5E). Therefore, the levels of extracellular L-Proline determine the outcome of cell competition both in cells and in the embryo. Collectively, these results indicate that during cell competition, defective cells increase L-Proline uptake from the extracellular environment. This increases intracellular L-Proline levels and induces their elimination by repressing the pro-survival ISR. This highlights how mitochondrial dysfunction sensitises cells to their metabolic environment and suggests that this is a mechanism that ensures that the epiblast is only comprised by the fittest cells during the early stages of post-implantation development.

## Discussion

Cell competition has been shown to regulate cell fitness in a wide range of contexts, from the developing embryo to the ageing tissue^3, 6, 26^. An important role of cell competition is to preserve tissue function by eliminating less-fit cells. A key feature of this process is that cells need to be able to measure and respond to the fitness levels of neighbouring cells. However, how a cell can sense the fitness of another is still poorly understood, with both mechanical and chemical signals potentially being involved^6^.

Here we have approached this problem by studying the fate of cells with mitochondrial dysfunction, the predominant type of defective cell eliminated by cell competition during early mouse development^17^. We find that during the first steps of pluripotent stem cell differentiation, mitochondrial dysfunction leads to activation of the ISR, a pathway that acts to restore cellular homeostasis in response to diverse stress stimuli^20, 21^. Both in the embryo and during pluripotent stem cell differentiation, these dysfunctional cells require the ISR for their survival, making them susceptible to repression of this pathway. Importantly, we show that in a heterogeneous environment, wild-type cells exploit this vulnerability and induce defective cell elimination via repression of the ISR and its key effector ATF4. This process allows for cells with robust mitochondrial activity to maximise their colonization of the tissue and potentially optimise tissue function.

Interestingly, we also find that when there is a high cell density, defective cells avoid elimination. This suggests that when high numbers of cells are damaged, activation of the ISR would allow to evade cell competition. Similar formation of a protective environment by increased numbers of loser cells has also been found to occur in the *Drosophila* imaginal wing disc^27^. The increased survival of defective cells as tissue crowding increases, argues that chemical signals rather than mechanical stresses likely determine the outcome of the cell competition occurring during differentiation.

But what chemical signals could regulate this cell competition? During the onset of differentiation, ESCs switch from relying on the uptake via macropinocytosism and digestion of exogenous proteins as a source of amino-acids, to the direct uptake of these metabolites^28^. Notably, exit of naïve pluripotency is required for competition in ESCs^15^. In consonance with these observations, we find a central role for the amino acid L-Proline in cell competition. In mouse ESCs, L-Proline has been suggested to play differing roles, from inducing proliferation of naïve cells and allowing these cells to adopt an early primed pluripotent state^29, 30^, to driving differentiation to a primitive ectoderm state^31, 32^. In contrast to these observations, we show that during the onset of differentiation, L-Proline induces the competitive elimination of cells with mitochondrial dysfunction. This is supported by several lines of evidence. First, in agreement with previous findings^25^, we identify that L-Proline represses the amino-acid starvation response branch of the ISR. Given that during differentiation the ISR is required for the survival of cells with mitochondrial dysfunction, this repression would be sufficient to induce dysfunctional cell elimination. Second, we find that in co-culture wild-type cells induce an increase in the expression of the enzymes that metabolise and import L-Proline in cells with mitochondrial dysfunction. This indicates that in a competitive environment, these dysfunctional cells will experience an increase in L-Proline metabolism, that correlates with the repression of the ISR that these cells show. Finally, we demonstrate that removing L-Proline, from the culture media of ESCs or mouse embryos is sufficient to restore ISR signalling in defective cells and prevents their elimination by cell competition. These results imply that rather than sensing the fitness of neighbouring cells, during cell competition wild-type cells overload defective cells with a metabolite that is toxic to them (Figure 6).

**Figure 6.**
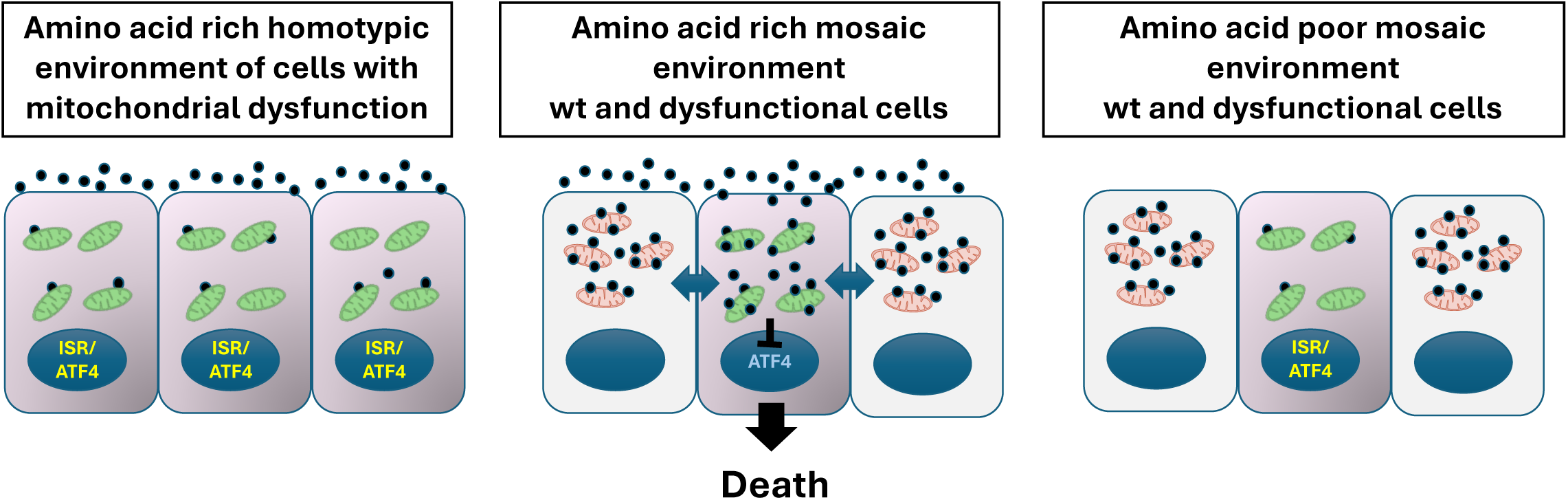
Model for defective cell elimination during cell competition. Diagrammatic representation showing how when cells with mitochondrial dysfunction are in a homotypic environment they have low intra-cellular levels of L-Proline (black circles) and ISR activation (left panel). In a competitive environment, the presence of wild-type cells induces increased L-Proline metabolism in the dysfunctional cells, repressing the ISR and causing their elimination (middle panel). In contrast to this, low levels of extra-cellular amino acids reduces L-Proline uptake in the cells with mitochondrial dysfunction, restoring ISR signaling and allowing for the survival (right panel).

The mechanism of cell competition that we describe here does not necessitate the sensing of relative fitness levels, but instead is caused by the relative susceptibility to extra-cellular metabolites. Two other pathways have been implicated in fitness sensing in *Drosophila* and in cancer cells: differential expression of FLOWER isoforms^33, 34^ and activation of the innate immune pathway^35^. It is interesting to note that viral infection, that can activate the innate immune pathway, is also a trigger for the ISR^20, 21^.. Similarly, FLOWER can act as calcium channel and changes in calcium levels can lead to unfolded proteins and ER stress, another trigger of the ISR. It will be interesting in the future to investigate any possible crosstalk between these pathways.

Finally, our results also have implications for the understanding of how nutrient availability in the extracellular environment shapes tissue composition and function. Our finding that L-Proline deficient media abolishes the negative selective pressure acting on defective cells, is counterintuitive as it suggests that the survival of dysfunctional cells is enhanced in nutrient poor conditions. Interestingly, this hypothesis is supported by one of the earliest findings of the cell competition field, that in fly, starvation prevents the competitive elimination of ribosomal mutant *Minute* clones from the *Drosophila* imaginal wing disc^36^. What these results imply is that cell competition optimises tissue function in response to nutrient availability, allowing for replacement of less fit cells to take place in nutrient rich environments but not in conditions of nutrient scarcity. Our work shows that amino acid sensing, and activation of the amino acid starvation response, plays a central role in the response to low nutrients, suggesting that protein levels in the diet could determine the nature of the competitive interactions occurring in the tissue. We propose that this would constitute a novel tissue sparing mechanism that would be protective for example in cases of maternal malnutrition or low protein diets. However, this protection would be at the expense of optimal tissue function, as it would allow the persistence of dysfunctional cell in the tissue.

## Acknowledgments

We would like to thank members of the Rodriguez lab for critical discussions and helpful suggestions, and Stephen Rothery for guidance and advice with confocal microscopy. We thank James Elliot, Joana De Teixeira Carrelha and Bhavik Patel from the LMS/NIHR Imperial Biomedical Research Centre Flow Cytometry Facility for support.

## Online Methods

### Cell lines and cell culture

E14, kindly provided by Prof A. Smith, from Cambridge University, were used as wild-type (WT) control cells tdTomato-labelled or unlabelled. GFP-labelled cells defective for BMP signalling (*Bmpr1a^-/-^*), cells null for Dynamin-related protein 1 (*Drp1^-/-^*) and *p53^-/-^*ESCs are described elsewhere^10, 15–17^. *Atf4^-/-^* and *p53^-/-^*; *Atf4^-/-^* ESCs were generated by CRISPR mutagenesis as previously described^16^.

Cells were cultured at 37 °C and 5% CO2 in GMEM-BHK21 supplemented with 10% FCS, 2mM glutamine, 1X MEM non-essential amino acids, 1mM sodium pyruvate, 0.1mM β mercaptoethanol (all Thermo Fisher Scientific) and 100 units/ml leukaemia inhibitory factor (generated and tested in the lab). The cells were grown in a monolayer, with the medium changed daily, and passaged every two days using Trypsin-EDTA (Sigma-Aldrich) for dissociation. Cell lines were routinely tested for mycoplasma contamination.

### ESC differentiation and cell competition assays

To induce differentiation, mESCs were seeded at low density (8x10^4^ cells/mL for 12-well plates, 1.6x10^5^ cells/mL for 6-well plates) on fibronectin-coated plates (Merck) in N2B27 media: an equimolar mix of DMEM/F-12 and Neurobasal media, containing 0.5X N2, 0.5X B27, 2mM L-glutamine, and 0.1mM 2-mercaptoetanol (all from Thermo Fisher). The media was changed daily, allowing differentiation to proceed for 3-4 days. All treatments began 24 hours after cell seeding and lasted for 48 hours. For L-proline treatment (+L-Proline), cells were cultured in N2B27 media with the addition of 0.5 mM L-proline. For halofuginone (Hg, Selleckchem) treatment, 20 nM of Hg dissolved in DMSO was supplemented to the N2B27 media. DMSO served as vehicle control. For pan caspase inhibitor treatment (Z-VAD-FMK, R&D Systems), differentiating cells were treated with 100uM of the inhibitor, with DMSO as vehicle control. L-proline deficient media (-L-Proline) was prepared by manually dissolving amino acids (Table 1), except L-proline, in water-based Neuronal Cell Culture Medium without Amino Acids (USBiological Life Sciences).

For cell competition assays, cells were seeded onto 12-well plates coated with fibronectin (Merck) at a density of 8x10^4^ cells per well for separate cultures and 4x10^4^ cells of each cell type at a 50:50 ratio per well for co-cultures. The cells were then cultured in differentiation medium. On day 3/4 of differentiation, cells were dissociated with Trypsin-EDTA, and cell counting was performed using the Countess™ 3 FL (Thermo Fisher), following the manufacturer’s instructions. Cells were resuspended in 3% FCS for fluorescence-activated cell sorting (FACS). The proportions of each cell type in co-cultures were determined using LSR II Flow Cytometer (BD Bioscience), based on the fluorescent tag of the ubiquitously expressed GFP or tdTomato in one of the cell populations.

### Mice and embryo culture

All mice were maintained on a 10 hr–14 hr light–dark cycle and treated in accordance with the Home Office’s Animals (Scientific Procedures) Act 1986. Wild-type mice analysed were maintained on a CD1 out-bred genetic background. Noon of the day of finding a vaginal plug was designated 0.5 dpc/. Embryo dissection was performed M2 media (Sigma). No distinction was made between male and female embryos during the analysis. For the caspase inhibitor, Hg and - L-Proline treatments, embryos were dissected at E5.5, cultured overnight (18 h) at 37 °C and 5% CO2 in 200 μM Z-VAD-FMK, 20nM Hg or the equivalent DMSO volume in N2B27 media (Neurobasal media; DMEM F12 media; 0.5X B27 supplement; 0.5X N-2 supplement; 0.1mM 2- mercaptoethanol; 2 mM glutamine (all Thermo Fisher Scientific), and fixed. For the -L-Proline experiments, embryos were cultured for 18h in either -L-Proline or control medium (N2B27 medium) and then fixed. After fixation, embryos were processed either for immunofluorescence analysis of cell death analysis by TUNEL staining. TUNEL staining was performed using *In Situ* Cell Death Detection Kit (Roche) following manufacturers protocol, as previously described^16^.

### Immunoblotting

Cells lysates were collected using RIPA buffer (50mM Tris- HCl at pH 6.8, 1%, 150mM sodium chloride, 1% NP-40, 1mM EDTA, 1X phosphatase inhibitor (Roche) and 1X protease inhibitor (Roche)), quantified using BCA quantification (Thermo Fisher Scientific) and resolved using Criterion XT pre-cast gels (BioRad) followed by transfer to PVDF membranes. Antibodies used are as follows: phospho-eIF2α, 1:1000, rabbit polyclonal cst 9721; ATF4, 1:1000, rabbit monoclonal, cst 11815; CHOP, 1:500, mouse monoclonal, sc 7351; cleaved CASPASE 3, 1:1000, rabbit polyclonal, cst 9661; PCNA, 1:5000, mouse monoclonal, sc56, β-ACTIN; 1:2500, rabbit polyclonal, cst 4967. Primary antibody incubation was performed overnight at 4 °C. This was followed by three TBST washes and secondary antibody incubation (anti-mouse (cst 7076) or anti-rabbit (cst 7074) conjugated to HRP; 1:4000) for 1.5 hours at room temperature. Blots were developed using ECL reagents (Merck) and imaged with the ChemiDoc™ MP system (Bio-Rad). To re-probe the membrane for alternative proteins, blots were stripped with stripping buffer (0.005% EDTA) by heating for 5 minutes. Protein expression analysis by densitometry was performed using ImageJ Fiji software and normalised to loading controls, β-ACTIN or PCNA.

### Immunofluorescence

Cells were fixed in 4% PFA for 10 min, permeabilised with 0.3% Triton X-100 (0.1% for cleaved CASPASE 3) for 5 min and blocked with 3% BSA (Sigma), 0.3% Triton X-100 (0.1% for cleaved caspase 3) for 1 h. Staining with primary antibodies (ATF4, 1:200, sc-390063 mouse monoclonal; CHOP, 1:200, cst 2895 mouse monoclonal; Cleaved CASPASE8; 1:100, cst 8592 rabbit monoclonal) were performed overnight at 4 °C in blocking buffer. After three washes in PBS, secondary antibodies (Alexafluor-647, Thermo Fisher Scientific; 1:500 dilution) and Hoescht (Thermo Fisher Scientific; 1:1000 dilution) were incubated for 1 h. Coverslips were then mounted on glass slides using Vectashield anti-fade mounting medium (Vector Labs).

Embryos were fixed using 4% PFA in PBS + 0.01% Triton X-100 and 0.1% Tween. Permeabilisation was performed with 0.5% Triton X-100 in PBS for 20 min and embryos were blocked using 2% horse serum in PBS + 0.1% Triton X-100 (PBT) for 45 min. Primary antibodies, as above, were incubated with embryos in blocking solution overnight at 4 °C. Following three washes in PBT, the embryos were incubated with secondary antibodies and DAPI, as above, for 1h at 4 °C. TUNEL staining was performed using *In Situ* Cell Death Detection Kit (Roche) following manufacturers protocol, as previously described^16^. Embryos were imaged in embryo dishes (Nunc) in a drop of Vectashield All images were captured on a Zeiss LSM-780 confocal microscope (40X oil objective lens-cells, 20X objective-embryos) and analysed using image j Fiji software^37^.

### Proliferation assays

Cells were incubated with EdU from the Click-iT kit (Thermo Fisher Scientific) for 2 hours according to manufacturer’s instructions. Cells were then detached from plates using accutase and analysed by flow cytometry using LSR II Flow Cytometer and FlowJo software.

### Bulk RNA-Seq

Cells grown for 3 days in N2B27 media were recovered into growth media and then resuspended in RLT lysis buffer (Qiagen). RNA extraction was performed using the Qiagen RNeasy kit according to the manufacturer’s instructions. Quality control, library preparation and sequencing were performed by the BRC Genomics Centre. RNA samples were quantified using a Qubit fluorometer (Thermo Fisher Scientific) and the quality assessed by TapeStation electrophoresis (Agilent). mRNA was isolated using oligo dT beads. mRNA was then fragmented, converted to cDNA and ligated to Illumina adapters. Following sample indexing, the quality of cDNA libraries was also assessed by TapeStation. Sequencing was performed using the Nextseq2000 system (Illumina). Differential gene expression analysis of the resulting sequencing data was performed by Nadia Fernandes (Imperial College London) in collaboration. Sequencing reads were aligned to the mouse genome (mm9) using TopHat2 (Kim et al. 2013) and differential expression was analysed using the DESeq2 package (Love et al. 2014). The enrichment analysis for the bulk RNA-seq datasets was performed using the g:Profiler tool76. The list of up-regulated, down- regulated and background genes related to the DE analysis for the bulk RNA-seq dataset are provided in the Supplementary Table 3.

### RNA extraction and quantitative RT-PCR

RNA was extracted with the RNeasy mini kit (Qiagen) and SuperScript III reverse transcriptase (Thermo Fisher Scientific) was used for cDNA synthesis according to manufacturer’s instructions. Quantitative RT-PCR was performed by amplification with Lightcycler 480 SYBR Green Master (Roche). The primers used are listed. RNA samples were collected from 3 independent experiments.

### Cell Sorting

E14 tdtomato cells and *Drp1^-/-^* cells were cultured as separate and co-cultures as mentioned above and then subjected to FACS-mediated cell sorting using BD FACSAria II sorter, followed by RNA extraction, cDNA synthesis and qPCR analysis. For FACS-mediated cell sorting, cells were trypsinised and resuspended in 500μl of 3%FBS in PBS. Cell suspensions were passed through a 40μm nylon mesh filter right before sorting. Post sorting, cells were collected in sort buffer (PBS, 1 mM EDTA, 25 mM HEPES pH 7.0, 1% FCS) and subjected immediately to RNA extraction as per method mentioned above.

### Metabolomic Analysis

For ESCs and differentiated samples, cells were cultured in -L-Proline medium supplemented with 0.21mM of labelled L-Proline (^13^C_5_-L-Proline (Cambridge Isotope Laboratories), kindly provided by Prof. Karen Vousden’s lab). Levels of labelled and unlabelled L-Proline were determined in cell culture supernatants and cell extracts.

At the time of harvest, medium samples were first collected from each well, and cells were washed with ice-cold PBS and scraped out in ice-cold HPLC-grade methanol containing 1nmol of nor- leucine (Merck, N8513) as an internal standard. The cell suspensions were collected in pre-chilled 2mL Eppendorf tubes and mixed with an equal volume of HPLC-grade chloroform and GCMS- grade water. The mixtures were then centrifuged at 14,800g for 10 minutes at 4°C to achieve biphasic partitioning. The polar metabolites in the upper phase of the samples were collected, . dried and resuspended in 100 μL methanol containing *scyllo*-Inositol (Merck, I8132) at 1 nmol per sample. The polar metabolite extracts were then transferred to a GCMS vial with insert. The samples were dried in a SpeedVac and washed twice with methanol.

For metabolite extraction from culture supernatants/medium, 5 μL of media per sample was added to a GCMS vial with insert. 10 uL 0.1mM *scyllo*-inositol (internal standard) was added to the vials, followed by drying in SpeedVac (∼15 minutes) and washing twice with methanol. Vials of 5 uL Metabolite Mix (MM) standards containing 1nmol of *scyllo*-inositol were prepared in separate vials as mentioned above.

Data acquisition was performed largely as previously described (PMID: 23763941) using an Agilent 7890B-5977A GC-MSD in EI mode after derivatization of twice methanol-washed dried polar extracts by addition of (a) 20 μL methoxyamine hydrochloride (20 mg/mL in pyridine (both Sigma) at room temperature overnight (>16 hours) and (b) 20 μL N,O- Bis(trimethylsilyl)trifluoroacetamide (BSTFA) containing 1% trimethylchlorosilane (TMCS) (Sigma) at room temperature for at least 1 hour. GC-MS parameters for polar analyses were as follows: carrier gas, helium; flow rate, 0.9 mL/min; column, DB-5MS (Agilent); inlet, 270°C; temperature gradient, 70°C (2 min), ramp to 295°C (12.5°C/min), ramp to 320°C (25°C/min, 3 min hold). Scan range was m/z 50-550 (polar). Data was acquired using MassHunter software (version B.07.02.1938). Data analysis was performed using MANIC software, an in house-developed adaptation of the GAVIN package (PMID: 21575589). Metabolites were identified and quantified by comparison to authentic standards, and label incorporation estimated as the percentage of the metabolite pool containing one or more ^13^C atoms after correction for natural abundance. In addition, we normalised peak areas to the cell number of the samples and internal standard *scyllo*- inositol.

### Quantification and Statistical Analysis

Flow cytometry data was analysed with FlowJo Software.

Western blot quantification was performed using Image software. Protein expression levels were normalised to loading controls β-actin or PCNA.

Statistical analysis was performed using GraphPad Prism v8.0.0 software. Statistical methods used are indicated in the relevant figure legends. For all *in vitro* analysis experiments, we used a sample size of 3 independent biological replicates, for ex vivo embryo experiments, we used a sample size of 8 biological replicates. Statistical significance was considered with a confidence interval of 0.05%; * p<0.05; ** p<0.01; *** p<0.005. Data obtained from cell competition assays, western blots and immunofluorescence quantification (multiple groups comparisons) were analyzed by two-way ANOVA, followed by Tukey’s multiple-comparison test. Data obtained from western blot, immunostaining analysis for embryos, proliferation assay were analysed by unpaired t-test.

#### Data Availability

Data were analysed with standard programs and packages, as detailed above. Authors can confirm that all relevant data are included in the paper and/ or its supplementary information files.

---

**Supplementary Figure 1.**
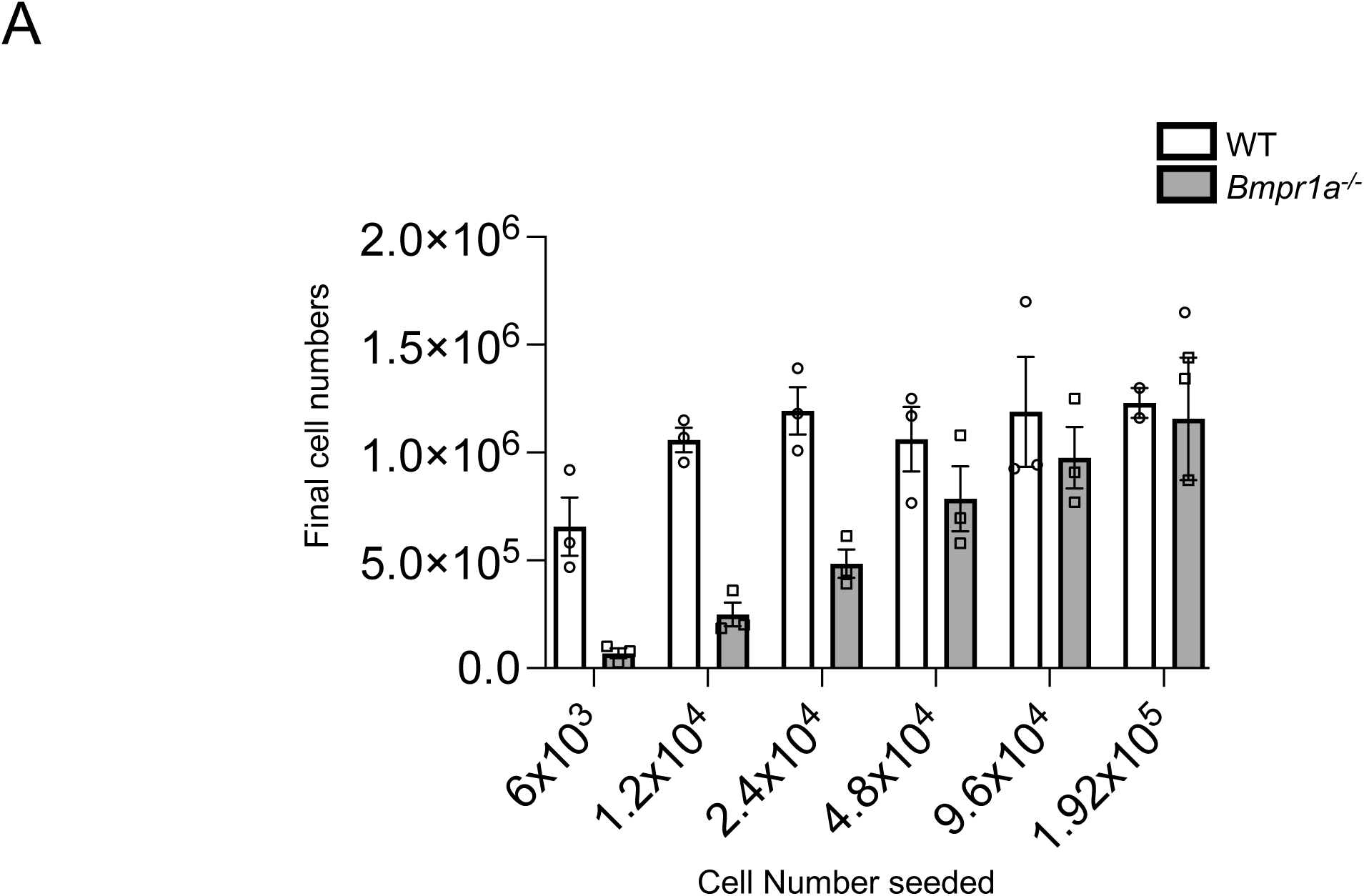
Bar graph representing final cell numbers of wild-type (WT) and *Bmpr1a^-/-^* cells when seeded at different cell densities. n=3. Error bars denote SEM. **p < 0.01, *p < 0.05, unpaired t-test.

**Supplementary Figure 2.**
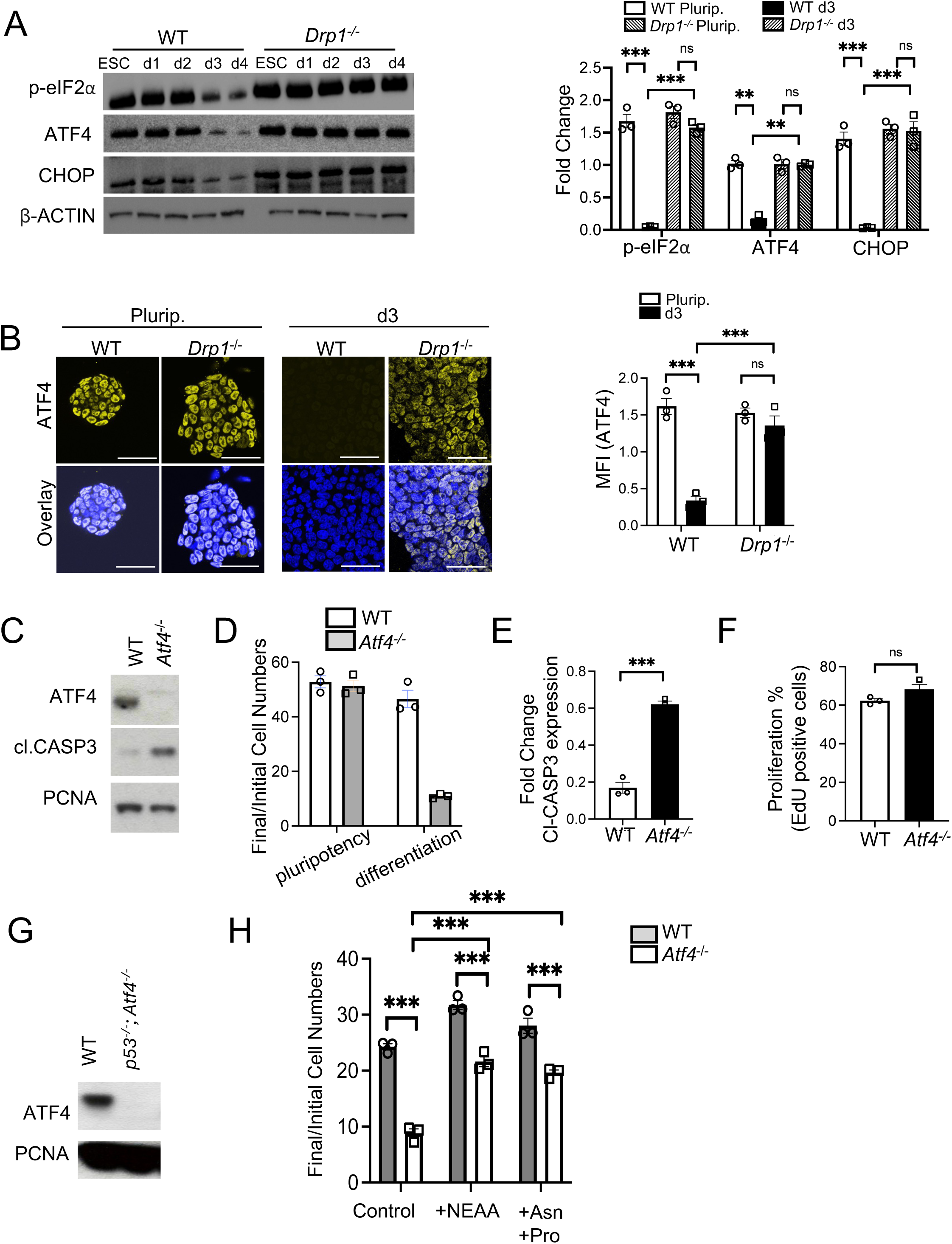
A, Wildtype (WT) and *Drp1^-/-^* cells were analysed by immunoblot after culture in pluripotency conditions (ESC) or at different time-points of differentiation (d1-d4) to detect expression levels of p-eIF2α, ATF4, CHOP and β-ACTIN (loading control). Bar graph represents fold change in expression levels of p-eIF2α, ATF4 and CHOP in ESC and d3 cells. B, Immunofluorescence analysis of ATF4 (yellow) in ESC and d3 WT and *Drp1^-/-^* cells, nuclei are counterstained with Hoechst (blue). Scale bars= 100uM. Bar graph represents quantification of staining intensity of ATF4 (MFI in arbitrary units) in the cells. C, immunoblot analysis showing ATF4 and cleaved-CASPASE 3 expression in WT and *Atf4^-/-^* cells. PCNA is used as a loading control. D, Bar graph depicting ratio of final (day 4) to initial cell numbers of WT and *Atf4^-/-^* cells cultured in pluripotency and differentiation conditions. E, Quantification of cleaved-CASPASE 3 levels from (C). F, Proliferation was assessed by EdU incorporation assay and bar graph depicts % of EdU incorporation in WT and *Atf4^-/-^* cells. G, immunoblot analysis showing ATF4 expression in WT and *p53^-/-^*; *Atf4^-/-^* cells. PCNA was used as a loading control. H, Bar graph representing ratio of final (d3) to initial cell numbers of WT and *Atf4^-/-^* cells cultured in control medium and in the presence of excess of non-essential amino acids (+NEAA) and excess of Asparagine and Proline (+Asn+Pro). n=3 for all studies. Error bars denote SEM. *** p < 0.005, **p < 0.01, *p < 0.05, ns- non-significant; two-way ANOVA and Tukeys post-hoc test (A, B, H, F). *** p < 0.005, ns-non- significant; unpaired t-test (D, E).

**Supplementary Figure 3.**
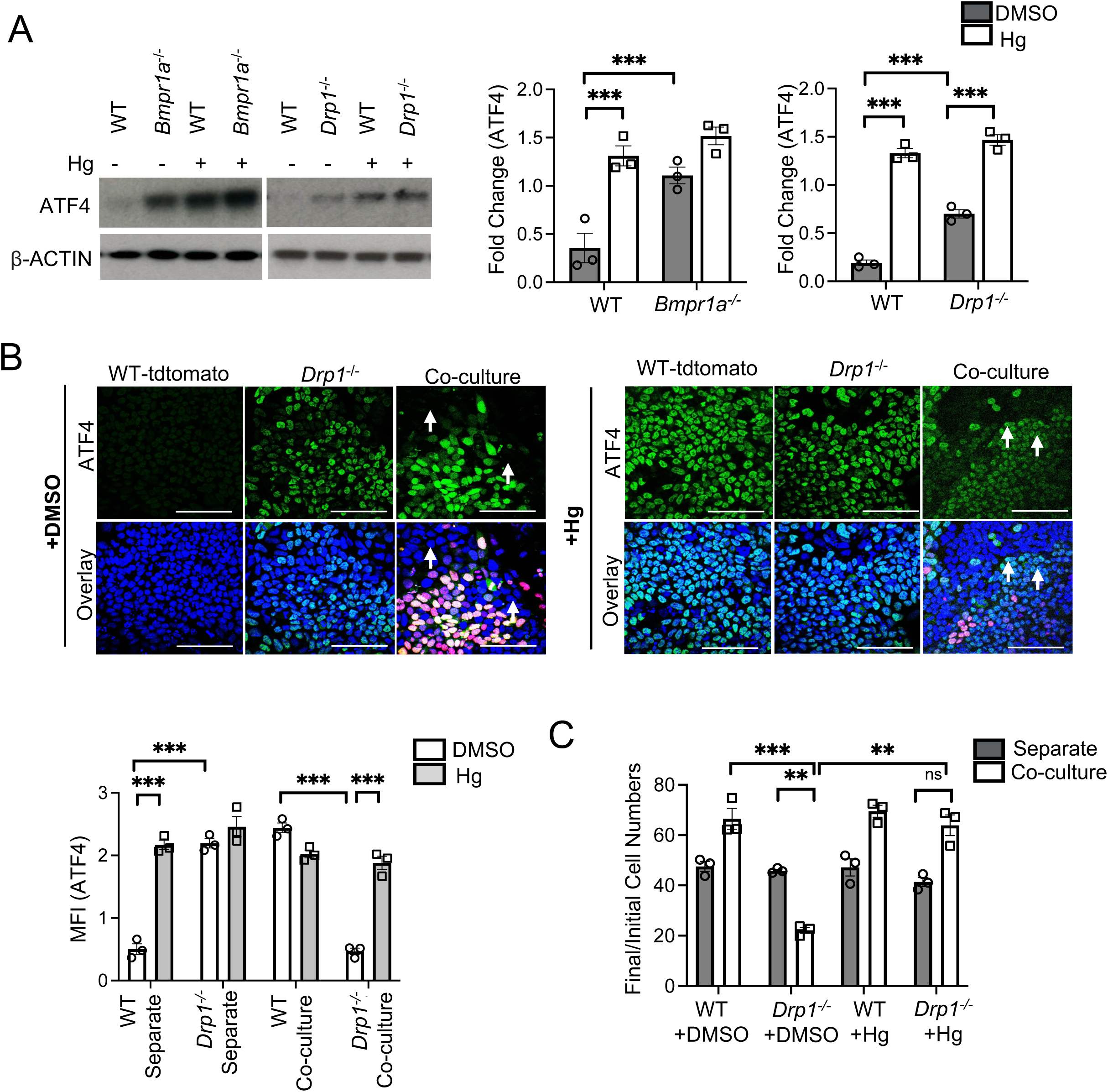
A, Wild-type (WT), *Bmpr1a^-/-^* and *Drp1^-/-^* cells were cultured in differentiation conditions and treated with DMSO or halofuginone (+Hg; 20nM) for 48h. Immunoblot analysis was performed to analyse expression levels of ATF4, β-ACTIN (loading control). Bar graphs represent densitometry-based quantification of ATF4 expression as fold change. B, Immunofluorescence analysis of ATF4 (green) in WT-tdtomato (red), *Drp1^-/-^* cells in separate and co-cultures after treatment with DMSO or Hg for 48h (d1-d3). Nuclei are counterstained with Hoechst (blue), white arrows indicate ATF4 expression levels in *Drp1^-/-^* cells in co-cultures. Scale bars= 100uM. Bar graph represents quantification of staining intensity of ATF4 (MFI in arbitrary units) in the cells. C, Cell competition assays between WT and *Drp1^-/-^* cells cultured for 48h with DMSO or Hg. The bar graph depicts the ratio of final cell numbers (day 3) to initial cell numbers for separate and co-cultures of these cells. n=3 for all studies. Error bars denote SEM. *** p < 0.005, **p < 0.01 ns-non-significant; two-way ANOVA and Tukeys post-hoc test.

**Supplementary Figure 4.**
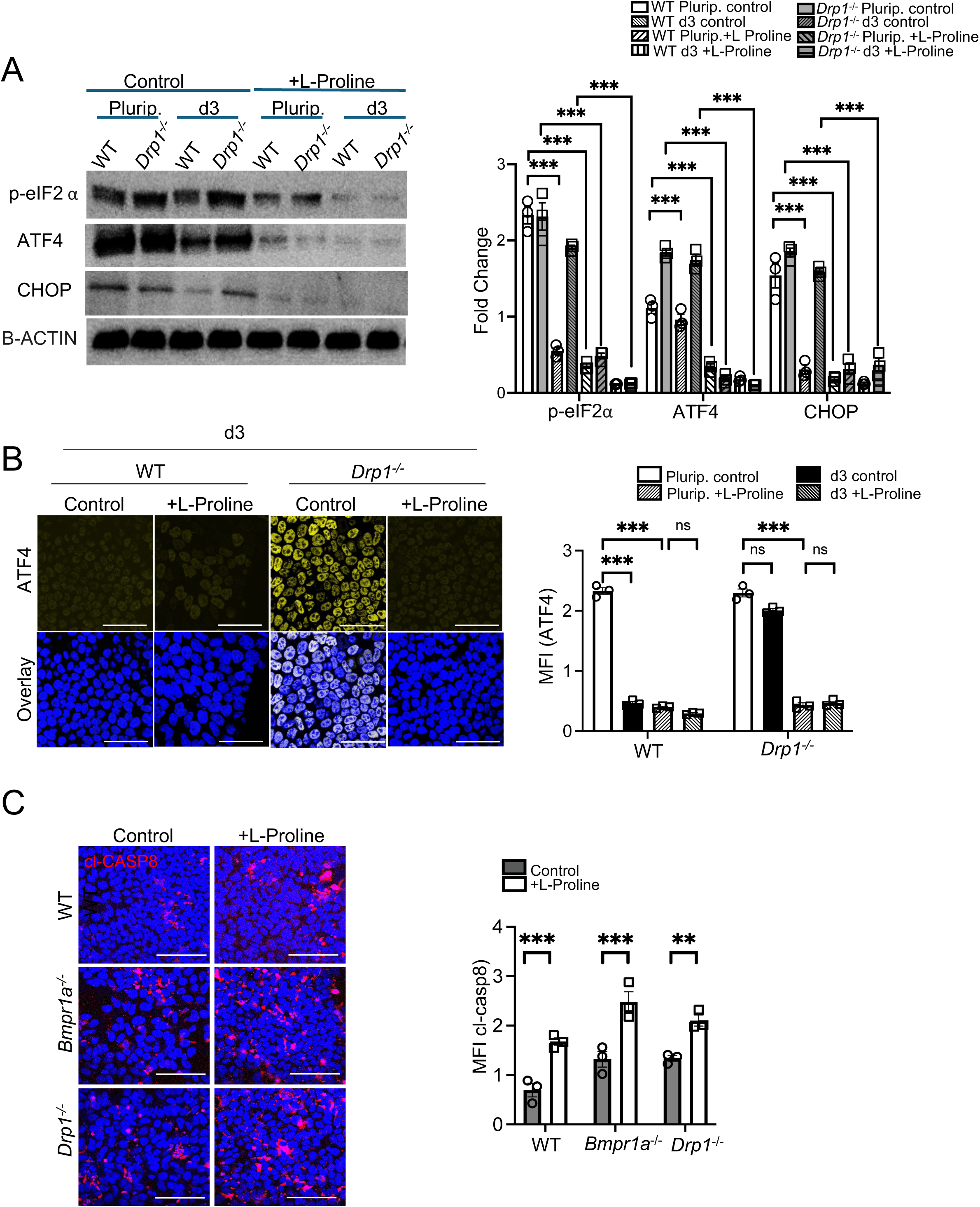
A, Wild-type (WT) and *Drp1^-/-^* cells were cultured in pluripotency (ESC) or differentiations (d3) conditions in control media or media supplemented with L-Proline for 48h (0.5mM; +L-Proline). Expression of ISR proteins, p-eIF2α, ATF4, CHOP was analysed by immunoblots. β-ACTIN was used as a loading control. Bar graph represents fold change in expression of the proteins in the various conditions. B, Immunofluorescence analysis of ATF4 in WT and *Drp1^-/-^* at day 3 of differentiation cultured in control medium or medium supplemented with L-Proline for 48h. Scale bar=100uM. Bar graph represents quantification of staining intensity of ATF4. C, Immunofluorescence analysis of cleaved caspase 8 (cl. CASP8) in WT *Bmpr1a^-/-^* and *Drp1^-/-^* cells cultured in control medium or medium supplemented with L-Proline for 48h. Scale bar=100uM. Bar graph represents quantification of staining intensity of cl. CASP8 (MFI in arbitrary units) in the cells. n=3 for all studies. Error bars denote SEM. *** p < 0.005, *p < 0.05 two-way ANOVA and Tukeys post-hoc test.

**Supplementary Figure 5.**
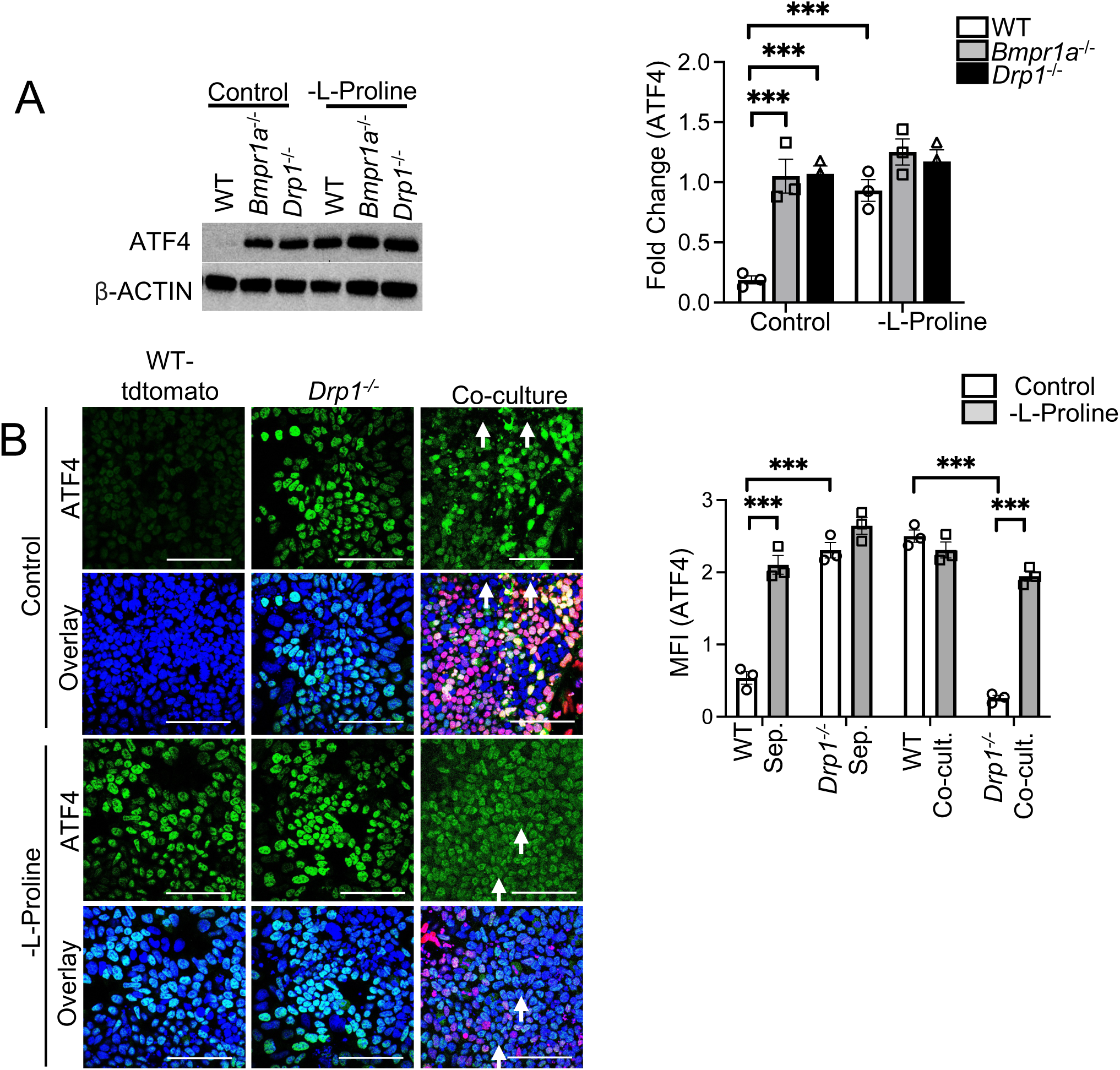
A, Wild-type (WT), *Bmpr1a^-/-^* and *Drp1^-/-^* cells were cultured in control differentiation medium and medium deprived of L-Proline (-L-Proline). Immunoblot analysis was performed to analyse expression levels of ATF4 and β-ACTIN (loading control) in these cells. Bar graphs represent densitometry-based quantification of ATF4 expression as a fold change. B, Immunofluorescence analysis of ATF4 (green) in WT-tdtomato (red), *Drp1^-/-^* cells in separate and co-cultures in control and -L-Proline medium. Nuclei are counterstained with Hoechst (blue), white arrows indicate ATF4 expression levels in *Drp1^-/-^* cells in co-cultures. Scale bars= 100uM. Bar graph represents quantification of staining intensity of ATF4 (MFI in arbitrary units) in the cells. n=3 for all studies. Error bars denote SEM. *** p < 0.005, *p < 0.05 two-way ANOVA and Tukeys post-hoc test.

**Supplementary Figure 6.**
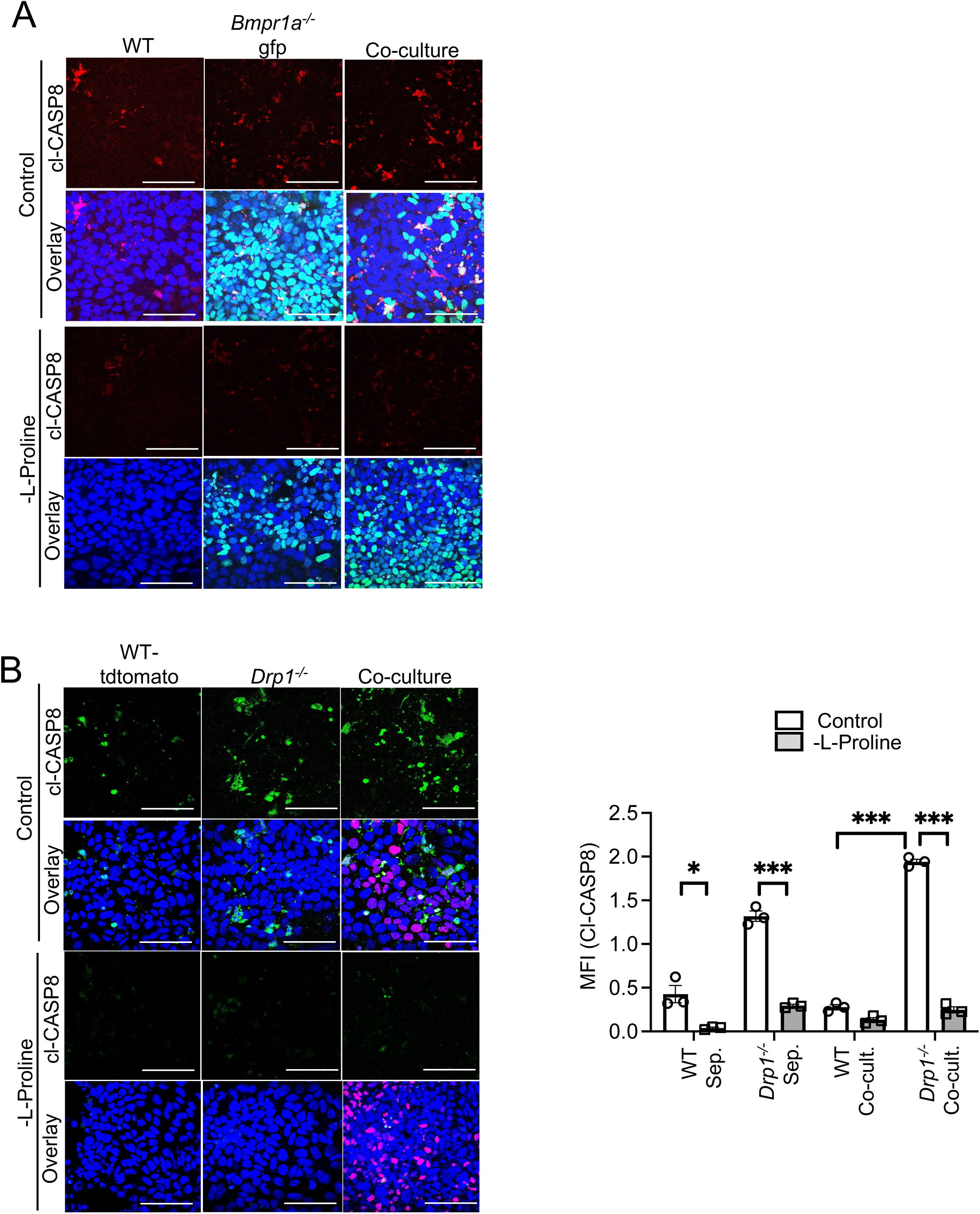
A, Immunofluorescence analysis of cleaved-CASPASE8 (red) in WT and *Bmpr1a^-/-^* (green) cells in separate and co-cultures in control and -L-Proline medium. Nuclei are counterstained with Hoechst (blue). Scale bars= 100uM. B, Immunofluorescence analysis of cleaved-CASPASE8 (green) in WT tdtomato (red) and *Drp1^-/-^* cells in separate and co-cultures in control and -L-Proline medium. Nuclei are counterstained with Hoechst (blue). Scale bars= 100uM. Bar graph represents quantification of staining intensity of cleaved CASPASE8 (MFI in arbitrary units) in the cells. n=3. Error bars denote SEM. *** p < 0.005, *p < 0.05 two-way ANOVA and Tukeys post-hoc test.

**Supplementary Figure 7.**
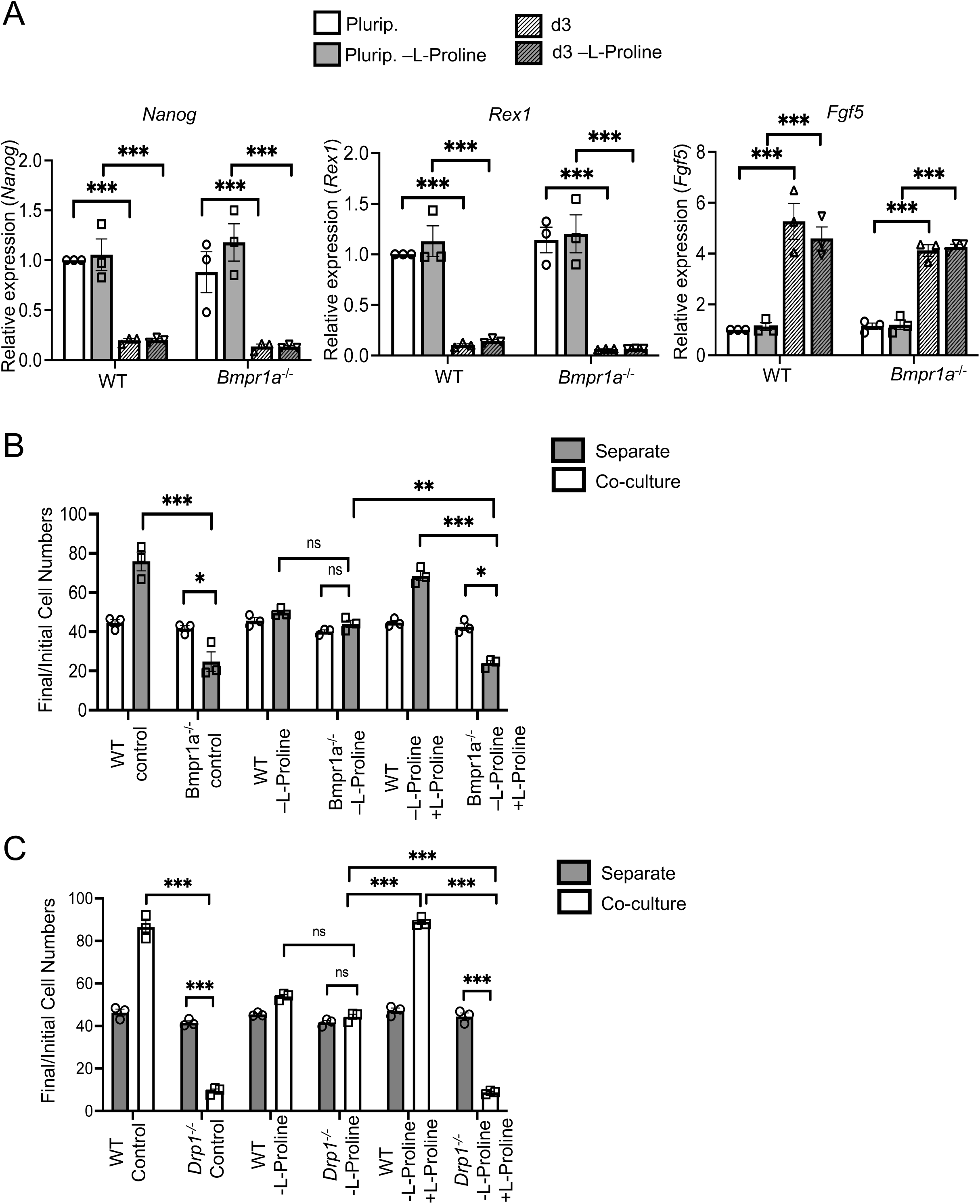
A, Quantitative RT-PCR showing gene expression levels of pluripotency (*Nanog* and *Rex1*) and differentiation markers (*Fgf5*) in wild-type (WT) and *Bmpr1a^- /-^* ESCs in pluripotency and at day 3 of differentiation. Gene expression is normalized against *Gapdh*. B, Cell competition assays between WT and *Bmpr1a^-/-^* cells in media lacking L-Proline with or without L-Proline treatment (0.25mM, 48h d1-d3), depicted as ratio of final cell numbers (day 3) to initial cell numbers for separate and co-cultures of these cells. C, Cell competition assays between WT and *Drp1^-/-^* cells lacking L-Proline with or without L-Proline treatment (0.25mM, 48h d1-d3), depicted as ratio of final cell numbers (day 3) to initial cell numbers for separate and co- cultures of these cells. n=3 for all studies. Error bars denote SEM. *** p < 0.005, *p < 0.05 two- way ANOVA and Tukeys post-hoc test.

**Supplementary Table 1:**
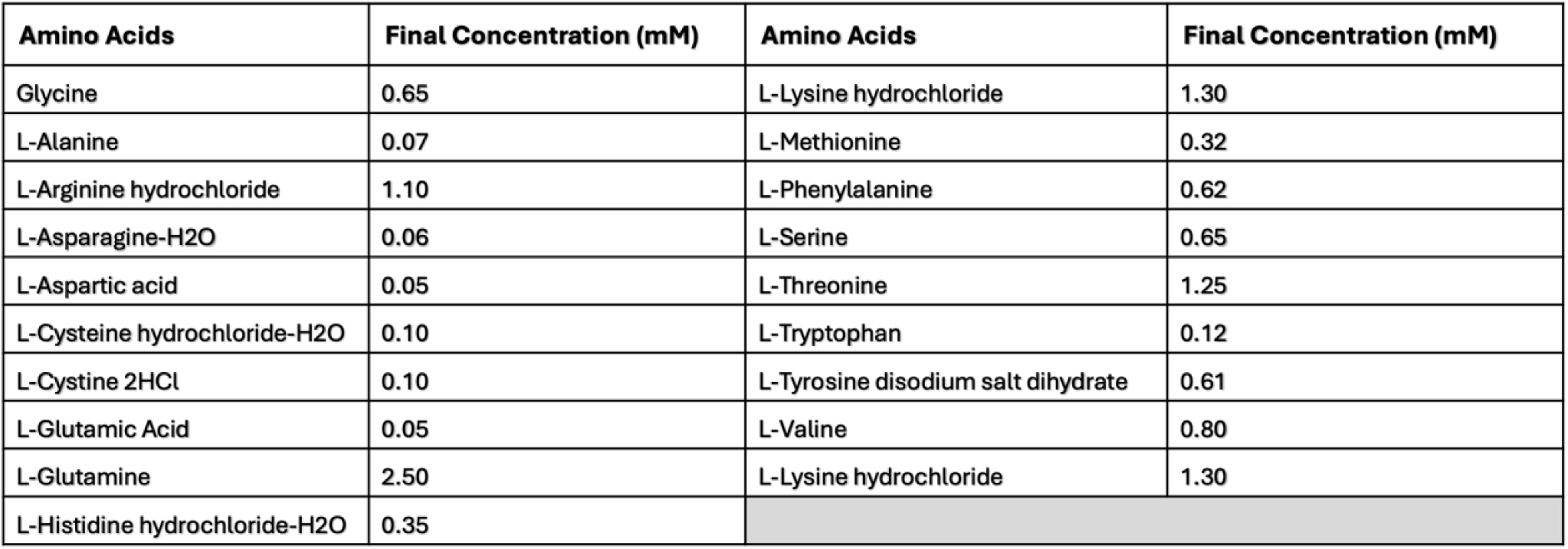
Composition of -L-Proline medium.

| Amino Acids | Final Concentration (mM) | Amino Acids | Final Concentration (mM) |
| --- | --- | --- | --- |
| Glycine | 0.65 | L-Lysine hydrochloride | 1.30 |
| L-Alanine | 0.07 | L-Methionine | 0.32 |
| L-Arginine hydrochloride | 1.10 | L-Phenylalanine | 0.62 |
| L-Asparagine-H<sub>2</sub>O | 0.06 | L-Serine | 0.65 |
| L-Aspartic acid | 0.05 | L-Threonine | 1.25 |
| L-Cysteine hydrochloride-H<sub>2</sub>O | 0.10 | L-Tryptophan | 0.12 |
| L-Cystine 2HCl | 0.10 | L-Tyrosine disodium salt dihydrate | 0.61 |
| L-Glutamic Acid | 0.05 | L-Valine | 0.80 |
| L-Glutamine | 2.50 | L-Lysine hydrochloride | 1.30 |
| L-Histidine hydrochloride-H<sub>2</sub>O | 0.35 |  |  |

**Supplementary Table 2:**
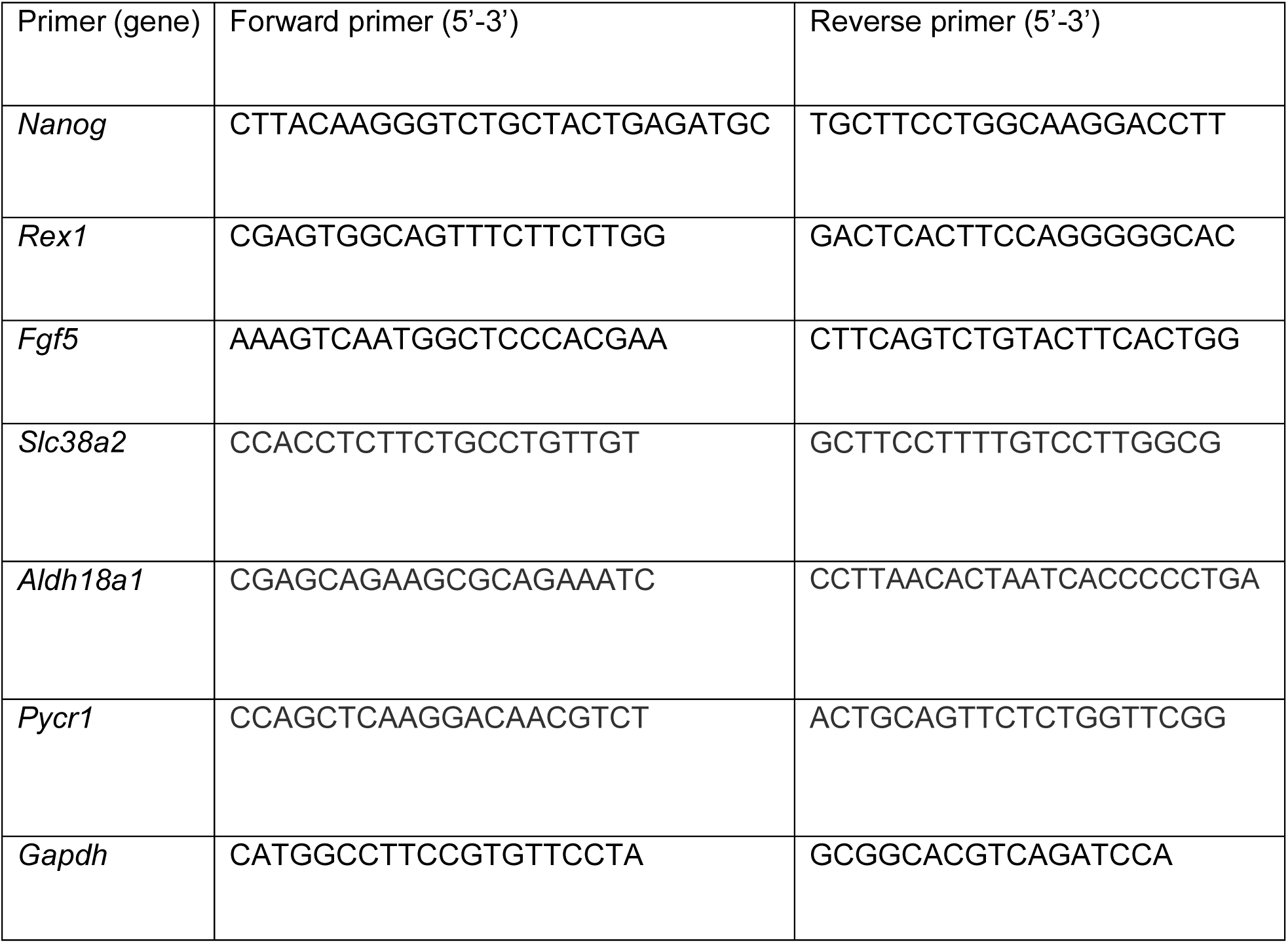
List of primers used for qPCR analysis.

## References

1. Morata, G. Cell competition: A historical perspective. Dev Biol 476, 33–40 (2021).

2. Nichols, J., Lima, A. & Rodriguez, T.A. Cell competition and the regulative nature of early mammalian development. Cell stem cell 29, 1018–1030 (2022).

3. Madan, E., Gogna, R. & Moreno, E. Cell competition in development: information from flies and vertebrates. Curr Opin Cell Biol 55, 150–157 (2018).

4. Esteban-Martinez, L. & Torres, M. Metabolic regulation of cell competition. Dev Biol. 475, 30–36 (2021).

5. Baker, N.E. Emerging mechanisms of cell competition. Nat Rev Genet 21, 683–697 (2020).

6. Bowling, S., Lawlor, K. & Rodriguez, T.A. Cell competition: the winners and losers of fitness selection. Development 146 (2019).

7. Claveria, C., Giovinazzo, G., Sierra, R. & Torres, M. Myc-driven endogenous cell competition in the early mammalian embryo. Nature 500, 39–44 (2013).

8. Moreno, E. & Basler, K. dMyc transforms cells into super-competitors. Cell 117, 117–129 (2004).

9. de la Cova, C., Abril, M., Bellosta, P., Gallant, P. & Johnston, L.A. Drosophila myc regulates organ size by inducing cell competition. Cell 117, 107–116 (2004).

10. Perez Montero, S., et al. Mutation of p53 increases the competitive ability of pluripotent stem cells. *Development (Cambridge*, England) 151 (2024).

11. Valverde-Lopez, J.A. et al. P53 and BCL-2 family proteins PUMA and NOXA define competitive fitness in pluripotent cell competition. PLoS Genet 20, e1011193 (2024).

12. Wagstaff, L. et al. Mechanical cell competition kills cells via induction of lethal p53 levels. Nat Commun 7, 11373 (2016).

13. Price, C.J. et al. Genetically variant human pluripotent stem cells selectively eliminate wild-type counterparts through YAP-mediated cell competition. Dev Cell (2021).

14. Vishwakarma, M. & Piddini, E. Outcompeting cancer. Nat Rev Cancer 20, 187–198 (2020).

15. Sancho, M. et al. Competitive interactions eliminate unfit embryonic stem cells at the onset of differentiation. Dev Cell 26, 19–30 (2013).

16. Bowling, S. et al. P53 and mTOR signalling determine fitness selection through cell competition during early mouse embryonic development. Nat Commun 9, 1763 (2018).

17. Lima, A. et al. Cell competition acts as a purifying selection to eliminate cells with mitochondrial defects during early mouse development. Nat Metab 3, 1091–1108 (2021).

18. Suomalainen, A. & Nunnari, J. Mitochondria at the crossroads of health and disease. Cell. 187, 2601–2627 (2024).

19. Lima, A., Burgstaller, J., Sanchez-Nieto, J.M. & Rodriguez, T.A. The Mitochondria and the Regulation of Cell Fitness During Early Mammalian Development. Curr Top Dev Biol 128, 339–363 (2018).

20. Pakos-Zebrucka, K. et al. The integrated stress response. EMBO Rep 17, 1374–1395 (2016).

21. Costa-Mattioli, M. & Walter, P. The integrated stress response: From mechanism to disease. Science 368 (2020).

22. Amiri, M. et al. Impact of eIF2alpha phosphorylation on the translational landscape of mouse embryonic stem cells. Cell Rep 43, 113615 (2024).

23. Labbe, K. et al. Specific activation of the integrated stress response uncovers regulation of central carbon metabolism and lipid droplet biogenesis. Nat Commun 15, 8301 (2024).

24. Keller, T.L. et al. Halofuginone and other febrifugine derivatives inhibit prolyl-tRNA synthetase. Nat Chem Biol 8, 311–317 (2012).

25. D’Aniello, C. et al. A novel autoregulatory loop between the Gcn2-Atf4 pathway and (L)- Proline [corrected] metabolism controls stem cell identity. Cell Death Differ 22, 1094–1105 (2015).

26. Diaz-Diaz, C. & Torres, M. Insights into the quantitative and dynamic aspects of Cell Competition. Curr Opin Cell Biol 60, 68–74 (2019).

27. Ballesteros-Arias, L., Saavedra, V. & Morata, G. Cell competition may function either as tumour-suppressing or as tumour-stimulating factor in Drosophila. Oncogene 33, 4377–4384 (2014).

28. Todorova, P.K. et al. Amino acid intake strategies define pluripotent cell states. Nat Metab 6, 127–140 (2024).

29. Casalino, L. et al. Control of embryonic stem cell metastability by L-proline catabolism. J Mol Cell Biol 3, 108–122 (2011).

30. Comes, S. et al. L-Proline induces a mesenchymal-like invasive program in embryonic stem cells by remodeling H3K9 and H3K36 methylation. Stem Cell Reports 1, 307–321 (2013).

31. Glover, H.J. et al. Signalling pathway crosstalk stimulated by L-proline drives mouse embryonic stem cells to primitive-ectoderm-like cells. *Development (Cambridge*, England) 150 (2023).

32. Washington, J.M. et al. L-Proline induces differentiation of ES cells: a novel role for an amino acid in the regulation of pluripotent cells in culture. Am J Physiol Cell Physiol 298, C982–992 (2010).

33. Rhiner, C. et al. Flower forms an extracellular code that reveals the fitness of a cell to its neighbors in Drosophila. Dev Cell 18, 985–998 (2010).

34. Madan, E. et al. Flower isoforms promote competitive growth in cancer. Nature 572, 260–264 (2019).

35. Meyer, S.N. et al. An ancient defense system eliminates unfit cells from developing tissues during cell competition. Science 346, 1258236 (2014).

36. Simpson, P. Parameters of Cell Competition in the Compartments of the Wing Disk of Drosophila. Developmental Biology 69, 182–193 (1979).

37. Schindelin, J., et al. Fiji: an open-source platform for biological-image analysis. Nat Methods 9, 676–682 (2012).

